# Conserved RNA–protein modules link early anthracycline responses to atrial fibrillation risk

**DOI:** 10.64898/2026.02.15.706019

**Authors:** Omar D. Johnson, E. Renee Matthews, Sayan Paul, José A. Gutiérrez, William K. Russell, Michelle C. Ward

## Abstract

Anthracycline chemotherapeutics increase risk for cardiovascular disease. The early molecular events that link inherited risk to cardiac dysfunction remain poorly defined. We therefore exposed iPSC-derived cardiomyocytes from six individuals to three anthracyclines and an anthracenedione, which act as topoisomerase inhibitors (TOP2i) to induce DNA damage, and generated proteomics data following three and 24 hours of TOP2i treatment. We constructed a 19 module co-expression network and identified four protein modules that associate with response to all TOP2i. Integration with a co-expression network generated from paired transcriptomic data revealed that the two most drug-responsive protein modules are preserved in the RNA network. The most preserved RNA module is enriched for p53 motifs in associated active regulatory regions and p53 target genes, which propagates to enrichment of p53 targets in the most drug-responsive preserved protein module. Integration of the protein network with genome-wide association studies across hundreds of cardiovascular traits identified a preserved protein module enriched for genetic risk of atrial fibrillation, PR interval, and longitudinal strain, suggesting that the molecular response to TOP2i is linked to genetic variation influencing cardiac electrophysiology and contractile performance. Differential protein abundance in this module associates with impaired calcium handling in cardiomyocytes thereby linking molecular effects to cellular decline. Together, these results define a TOP2i-induced gene regulatory response propagated to the proteome that underlies early cardiotoxicity, and demonstrate how a multi-level network architecture can provide insight into genetically mediated susceptibility to drug-induced stress.

## INTRODUCTION

Chemotherapy-related cardiac dysfunction (CTRCD) has become a leading cause of non-cancer morbidity among cancer survivors (Gilchrist et al. 2019; Sturgeon et al. 2019). Up to 10% of patients treated with anthracycline (AC) drugs develop symptomatic heart failure within five years, and many more show subclinical left-ventricular dysfunction (Denlinger et al. 2018; Bostany et al. 2025). AC treatment also increases the risk of atrial fibrillation, which can exacerbate left-ventricular dysfunction and elevate stroke risk (McGowan et al. 2017; Cardinale et al. 2020; Fradley et al. 2021). Clinical identification of CTRCD depends on late markers, such as increased circulation of cardiac troponin I or a significant decrease in the left-ventricular ejection fraction, by which time myocardial injury is already established (Park et al. 2017; Yan et al. 2020; Canty Jr 2022). Speckle-tracking echocardiography can detect early impairment of longitudinal strain (LS), a measurement of myocardial contractility that precedes other clinical indications of CTRCD; however it does not offer insight into the molecular events that are coupled to early functional decline (Thavendiranathan et al. 2014; Camilli et al. 2024). To identify targets to prevent or reverse the effects of CTRCD, it is therefore critical to determine the molecular signatures coupled to early contractile impairment.

Topoisomerase II (TOP2) inhibitors (TOP2i), including ACs such as Doxorubicin (DOX), Daunorubicin (DAU) and Epirubicin (EPI), as well as the anthracenedione Mitoxantrone (MITO), induce cancer cell death by trapping TOP2 on DNA leading to the generation of DNA double-strand breaks (Yang et al. 2014). In cancer cells TOP2i-induced DNA double-strand breaks are mediated by the α-isoform (TOP2α) that is highly expressed in proliferating cells. Terminally-differentiated cardiomyocytes express high levels of the β-isoform of TOP2 (TOP2β) rendering these non-cancerous cells especially vulnerable to TOP2-linked DNA damage. The only FDA-approved drug designed to reduce the risk of DOX-induced cardiotoxicity is dexrazoxane, which degrades TOP2β. While dexrazoxane affords partial myocardial protection from ACs at high cumulative doses, concerns about attenuated anticancer efficacy and secondary malignancies have constrained its use, leading to a search for alternatives (Tebbi et al. 2007; Burridge et al. 2016; Jirkovský et al. 2021; Arzt et al. 2023). A landmark study by Liu *et al*. screened 2,300 genes in induced pluripotent stem-cell–derived cardiomyocytes (iPSC-CMs) and identified carbonic anhydrase 12 (CA12) as a therapeutic target to mitigate DOX-induced cardiotoxicity, thereby offering an alternative to dexrazoxane (Liu et al. 2024). However, this study restricted screening to genes encoding currently druggable proteins. Given the rapid advancement of therapeutic modalities across molecular phenotypes, there is a pressing need for unbiased approaches to identify candidate gene sets associated with CTRCD to enable broader functional follow-up.

IPSC-CMs treated with ACs recapitulate many features of CTRCD and enable highly controlled mechanistic studies of the transition between healthy and disease states (Burridge et al. 2016; Adegunsoye et al. 2023). Transcriptomic profiling of iPSC-CMs has identified thousands of differentially expressed genes associated with calcium dysregulation within hours of clinically relevant sub-micromolar, and sub-lethal treatment of TOP2i (Knowles et al. 2018; Matthews et al. 2024). Proteomic data, however, remain sparse (Johnson et al. 2025). It is therefore unknown whether these early transcriptional changes propagate to the proteome across TOP2i drugs. This is particularly important to understand given that protein abundance is more directly linked to cellular function than mRNA levels due to post-transcriptional, translational, and degradation-based regulation (Vogel and Marcotte 2012; Lee and Yaffe 2016; Liu et al. 2016; Munro et al. 2024). It is therefore unclear how molecular phenotypes coordinate in response to TOP2i treatment in human cardiomyocytes and connect to functional and genetic indicators of cardiotoxic risk.

Here, we generated quantitative proteomic profiles from iPSC-CMs derived from six individuals treated with sub-lethal, clinically relevant concentrations of DOX, EPI, DAU, and MITO for three and 24 hours, and integrated these with paired transcriptomic data (Matthews et al. 2024). We constructed RNA and protein co-expression networks to identify drug-specific and shared responses, enabling us to capture time-resolved, multi-layered molecular phenotypes to link to cardiovascular traits and disease.

## RESULTS

We designed a study to measure the response of human cardiomyocytes to TOP2i chemotherapeutic agents at the level of the transcriptome and proteome (Fig 1A). We differentiated iPSCs from six healthy female individuals with no known history of disease into iPSC-CMs (See Methods). iPSC-CM cultures contained a high proportion of cardiac troponin T expressing cells across individuals indicating successful cardiac differentiation (median purity = 95.1%; See Methods; Table S1).

**Figure 1:**
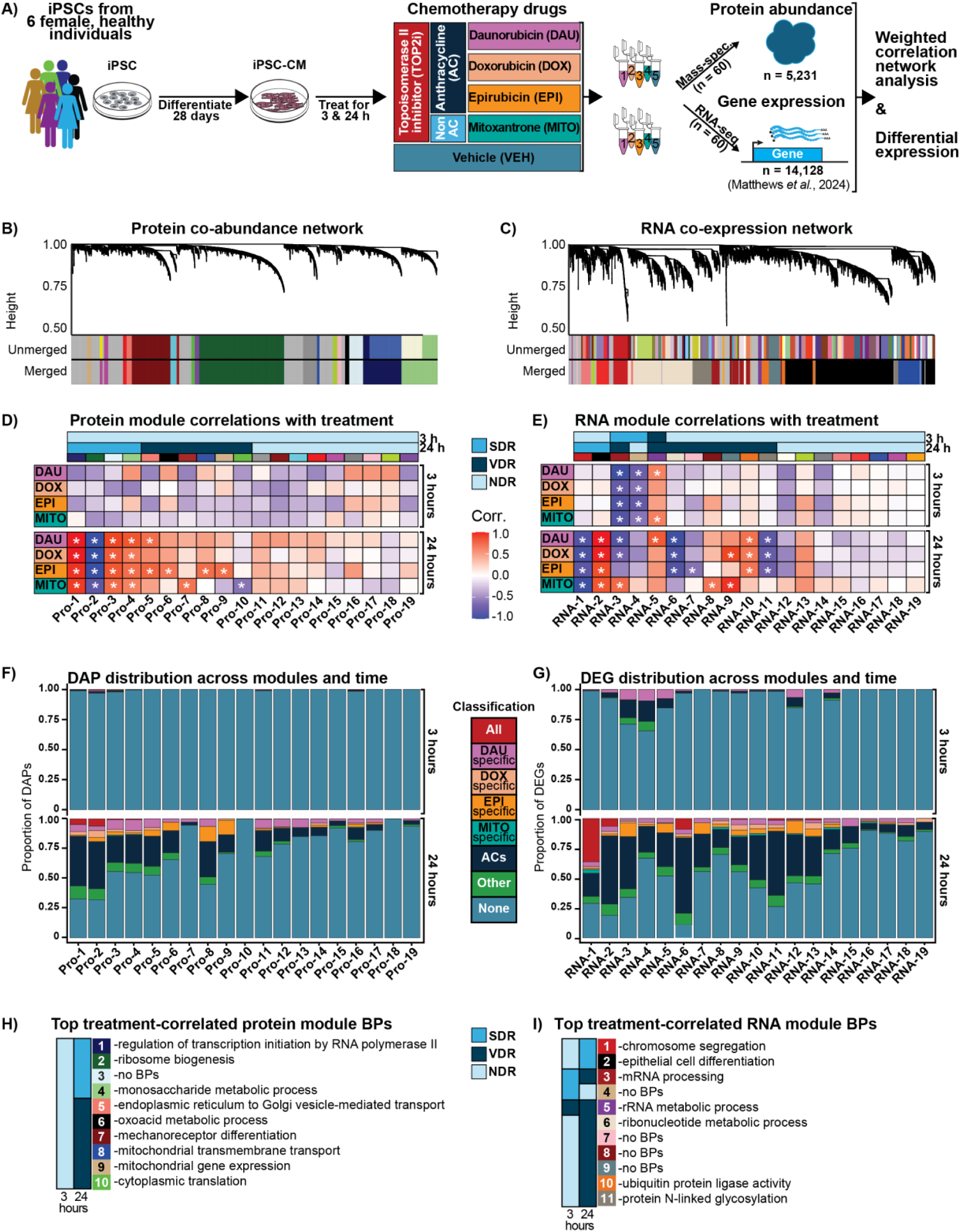
TOP2i treatment in iPSC-CMs induces RNA and protein co-expression changes in a time- and drug-dependent manner. **A)** Experimental design of the study. iPSCs derived from six healthy females aged 21 to 32 were differentiated into cardiomyocytes (iPSC-CMs) and treated with a panel of TOP2i drugs used in cancer treatment. The response to Daunorubicin (DAU), Doxorubicin (DOX), Epirubicin (EPI), and Mitoxantrone (MITO) was compared to a water vehicle (VEH) at three and 24 hours post treatment. 60 experimental samples from the six individuals were used in mass spectrometry to measure protein abundance and 60 matched samples were used to measure gene expression by RNA-seq (Matthews et al. 2024). **B)** Cluster dendrogram of the protein co-abundance network. Height represents the dissimilarity of clusters across modules. Each module is shown by a different color prior to and after merging of related modules. **C)** Cluster dendrogram of the RNA co-expression network. Each network is generated independently of the other. **D)** The correlation between each eigenprotein and drug treatment in the protein network. Asterisk represents a significant correlation between the eigenprotein and the trait (\**P* < 0.05). Modules are ordered from highest absolute median correlation across all drug treatments and time to lowest. Module designations for Shared Drug Response (SDR), Variable Drug Response (VDR) and No Drug Response (NDR) at either three or 24 hours is included above the correlations. **E)** The correlation between each eigengene and drug treatment in the RNA network, following the same convention as D. **F)** Distribution of three- and 24-hour differentially abundant proteins (DAPs) across protein modules. Colors designate DAPs shared in all drug treatments (All), specifically in one drug treatment, and shared across anthracyclines (AC). DAPs identified in two or three drug treatments that are not contained in the above-mentioned categories are represented as ‘Other’, and proteins that are not DAPs in response to any drug treatment are designated as ‘None’. **G)** Distribution of three- and 24-hour differentially expressed genes (DEGs) across RNA modules. DEGs are further classified as described for DAPs. **H)** Top enriched biological process (BP) in each protein module (Fisher exact test; adj. *P* < 0.05). If a module is not enriched for a BP then it is designated as ‘no BPs’. **H)** Top enriched BP in each RNA module.

Our previous work indicated that calcium handling in iPSC-CMs is affected within 24 hours of TOP2i treatment at a concentration that is clinically relevant and sub-lethal (0.5 μM; below the LD50) (Matthews et al. 2024). We first verified that 24 hours of treatment with 0.5 μM of three ACs DAU, DOX and EPI and one anthracenedione MITO does not affect iPSC-CM viability when compared to a water vehicle (VEH) control (Fig S1). To systematically examine the effects of TOP2i on cardiomyocytes across molecular phenotypes, we therefore treated iPSC-CMs from each individual with 0.5 μM DAU, DOX, EPI MITO and VEH. We selected three and 24 hour treatment times given the range of transcriptional responses across these timepoints (Matthews et al. 2024). Cells from the same differentiation and treatment batch were collected for both RNA and protein extraction, yielding 120 total experimental samples for downstream analysis (See Methods; Table S1).

Given the large sample number, we carefully designed treatment- and time-balanced batches for RNA and protein extractions and high-throughput analysis (Table S1). The RNA data is reported in Matthews *et al*. (Matthews et al. 2024). The 60 sample protein data was generated by extracting protein from iPSC-CMs in six batches corresponding to all treatments and timepoints per individual (n = 10). Protein samples were organized for digestion in three randomized blocks containing all samples from two individuals, and proteomics analysis was performed in data-independent acquisition (DIA) mode in six batches, one for each individual, with block randomization for each batch. To account for machine drift over time, a pooled digest of the 10 samples from each block was run before and after all randomized samples, along with a pooled digest of all 60 samples at the end of each run. Analysis of these pools indicated minimal variability in protein abundances due to drift (Fig S2). We combined the 60 experimental samples with the 60-sample pool analyzed in every mass spectrometry batch to estimate and remove latent technical variation. After correction for unwanted factors, pooled samples were removed from the data set.

We defined protein expression measurements as the protein abundance data from the 60 unique protein samples (S3 Appendix), and gene expression as the log_2_ cpm-standarized counts from the 60 RNA samples (Matthews et al. 2024). After filtering out proteins with missing or low abundance values and RNA with low expression levels, we retained 5,231 proteins and 14,128 genes for downstream analysis (See Methods). Principal component analysis (PCA) reveals that PC1 accounts for 20.27% of the variation in protein abundance and 41.2% of the variance in gene expression, and associates with both treatment type and treatment time across phenotypes (Fig S3).

### Network analysis identifies co-expressed RNA modules and co-abundant protein modules

To facilitate a systems-level understanding of the drug- and time-dependent molecular responses to TOP2i, we applied Weighted Gene Co-expression Network Analysis (WGCNA) independently to the RNA and protein datasets. Networks were optimized to reflect a scale-free topology. Specifically, we systematically evaluated connectivity frequency distributions across 20 soft-thresholding power values, selecting thresholds that achieved high scale-free topology fits (R^2^ between 0.8 - 0.9). These thresholds were chosen based on established guidelines to balance the inclusion of biologically meaningful connections while minimizing the incorporation of spurious associations (See Methods) (Langfelder and Horvath 2008). We subsequently minimized module redundancy within each network by merging highly similar modules based on eigenprotein and eigengene correlations, followed by Gene Ontology (GO) enrichment analyses to confirm functional distinctness of module identities. In the protein co-abundance network we selected a soft power of 10 (R^2^ = 0.83; Fig S4A), which resulted in a mean connectivity score of 34.7, and minimized module redundancy via merging of functionally-related and highly-correlated modules (r > 0.8; Pearson correlation; Fig 1B). This approach yields 18 co-abundant protein modules (Table S2), each represented by an eigenprotein (Fig S4B), and one module containing proteins that are not co-abundant (Pro-16; See Methods). Each module contains between 35 - 1,257 proteins (Fig S4C). For the RNA co-expression network, we selected a soft power of 16 (R^2^ = 0.9; Fig S5A), which resulted in a mean connectivity score of 102, and minimized module redundancy via merging of functionally-related and highly-correlated modules (r > 0.7; Fig 1C). This produced 18 co-expressed RNA modules (Fig S5B; Table S3), and one module containing genes that are not co-expressed (RNA-14). Each RNA module contains between 48 - 4,523 genes (Fig S5C).

A characteristic feature of scale-free networks is the presence of a small number of highly connected hubs that are essential for maintaining network stability and represent module properties. We identify 388 hub proteins and 1,239 hub genes that are highly connected (within the top 10% by intra-modular connectivity (kIN)) that have a strong correlation with the respective module eigengene or eigenprotein (kME > 0.7).

### RNA and protein modules correlate with TOP2i treatment in a drug- and time-dependent manner

To associate protein and RNA modules with response to drug treatment, we calculated pairwise correlations between the eigenvalues of drug-treated samples and their time-matched VEH controls at either the three- or 24-hour timepoint for each module. Modules were ranked based on their absolute median drug correlation strength. Ten protein modules (Pro-1 to Pro-10) are correlated with at least one drug 24 hours following treatment (*P* < 0.05), while no protein modules are correlated with any drug after three hours (Fig 1D). We further classified modules based on whether they are correlated with all drugs at a given timepoint (Shared Drug Response; SDR; Pro-1 to Pro-4), at least one drug (Variable Drug Response; VDR; Pro-5 to Pro-10), or no drugs (No Drug Response; NDR; Pro-11 to Pro-19). Pro-1, Pro-3, and Pro-4 SDR modules are positively correlated with all treatments indicating that protein abundance increases in response to treatment, while Pro-2 is negatively correlated. Pro-5 is positively correlated with DAU and EPI. Pro-6, Pro-8, and Pro-9 are positively correlated with EPI only, while MITO correlates only with Pro-7 (positively) and Pro-10 (negatively).

The RNA co-expression network includes eleven RNA modules (RNA-1 to RNA-11) that are correlated with at least one drug and ordered based on absolute median correlation to all drugs (Fig 1E). In contrast to the protein network, three of these modules are correlated with at least one drug following three hours of treatment. RNA-1 is a negatively correlated SDR module at 24 hours and RNA-2 a positively correlated SDR module at 24 hours. Neither module is correlated with any drug at three hours. RNA-3 and RNA-4 are negatively correlated SDR modules at three hours but have differing effects at 24 hours - RNA-3 is a VDR module while RNA-4 is an NDR module. RNA-5 is a VDR module at both timepoints, while RNA-6 to RNA-11 are VDR modules at 24 hours but NDR modules at three hours. RNA-6, RNA-10 and RNA-11 are correlated with all ACs but not MTX. RNA-12 to RNA-19 are denoted NDR at both timepoints. These findings show that protein and RNA co-expression responses are both time- and drug-dependent, and that protein responses emerge later in time than RNA responses.

### Shared drug response modules are enriched in differentially expressed genes and proteins

We independently determined the protein and RNA responses to TOP2i treatment by performing pairwise differential expression (RNA) and pairwise differential abundance (protein) testing for each drug relative to its time-matched VEH control at the three- and 24-hour timepoints (See Methods). At three hours, 46 differentially abundant proteins (DAPs; adjusted *P* < 0.05) and 505 differentially expressed genes (DEGs; adjusted *P* < 0.05) are identified in response to at least one drug (Tables S4-S11), whereas at 24 hours, 1,619 DAPs and 7,857 DEGs are detected (Tables S12-19). ACs induce more expression changes at both the protein and RNA level compared to MTX following 24 hours of treatment. 19-23% of proteins are DAPs in response to each AC (n = 973-1,225) compared to 4% (n = 202) of proteins that are DAPs in response to MTX, and 45-50% of genes are categorized as DEGs in response to each AC compared to 8% in response to MTX (Fig S6).

We asked whether the patterns of drug correlation across co-expression modules aligns with the proteins and genes that were determined to be differentially expressed in response to one or more drug treatments. We therefore assessed the proportion of DAPs and DEGs in each protein and RNA network module. In line with the lack of any drug correlation with any protein module following three hours of treatment, the proportion of three-hour DAPs in three-hour samples in every module is low (< 3%; Fig 1F). However, at 24 hours, SDR modules contain up to 60% DAPs. Drug-correlated modules exhibit a higher proportion of DAPs than non-drug-correlated modules (median SDR+VDR = 31%, median NDR = 7.7%). Among drug-correlated modules, DAPs shared across all ACs make up the largest fraction of module DAPs (0 - 52.2%; median = 28.5%). SDR modules Pro-1, Pro-2, Pro-3 and Pro-4 contain 1-7% DAPs shared across all drug treatments, with Pro-1 and Pro-2 showing the greatest proportion (6-7%), aligning with their SDR classification at this timepoint. The three-hour drug treatment samples in the RNA network show highest enrichment for DEGs in the SDR modules (up to 20% of module genes; Fig 1G); while the 24-hour samples show up to 87% of SDR module genes classified as DEGs. Drug-correlated modules show a higher proportion of DEGs than non-drug-correlated modules (median SDR+VDR = 48%, median NDR = 13%). DEGs shared across all ACs comprise the largest fraction of DEGs in all drug-correlated modules except RNA-1, which is predominantly composed of DEGs shared across all drugs (38.5%). These data indicate that protein and RNA co-expression modules comprise proteins and genes that respond to drug treatments in line with the module level drug correlation patterns, and that there is a high degree of sharing in the drug response across TOP2i drugs and ACs specifically.

### Drug-correlated modules associate with distinct functions

None of the TOP2i affect protein levels of cardiac troponin T and cardiac troponin I, the clinically-measurable markers of cardiac damage. However, TOP2i are known to affect TOP2β function in cardiomyocytes, and activate the TP53 DNA damage response. We therefore investigated the TOP2β and TP53 response to treatment and localization within the RNA and protein co-expression networks. TOP2β is classified as both a downregulated DAP and DEG in response to treatment with all drugs following 24 hours of treatment, and is localized in SDR module Pro-2 and VDR module RNA-6 (Fig S7). TP53 is a hub protein in Pro-1 and is upregulated and differentially abundant in all 24-hour drug treatments but is neither a DEG nor a hub gene. These findings demonstrate that proteins canonically associated with TOP2i mechanisms are consistently altered across treatments. Integration between the RNA and protein networks provides insights into the molecular level at which effects on protein originate. Our framework also enables the identification of novel proteins critical to the TOP2i response. For instance, POLR2A, FCF1 and hub protein TBL3 are DAPs in the Pro-2 module and are downregulated in response to all drug treatments at both three and 24 hours; however their RNA levels are variably affected by the different drugs over time suggesting molecular phenotype-specific effects (Fig S8). These three proteins relate to transcription and rRNA processing, suggesting that these processes are commonly disrupted in response to TOP2i treatment.

To understand broad effects of these drugs on cellular pathways, we tested for enrichment of biological processes in modules associated with drug treatment (SDR or VDR) relative to all detected proteins or expressed genes using gene ontology analysis (Fisher’s exact; adjusted *P* < 0.05). Of the 10 drug-correlated protein modules, nine are enriched for at least one biological process (BP) (Fig 1H), while seven of 11 drug-correlated RNA modules show BP enrichment (Fig 1I). Among protein SDR modules, which represent core treatment responses across drugs, key enriched processes include regulation of transcription initiation by RNA polymerase II (Pro-1), ribosome biogenesis (Pro-2), and monosaccharide metabolism (Pro-4). In the RNA network, SDR modules are enriched for chromosome segregation (RNA-1), epithelial cell differentiation (RNA-2), and mRNA processing (RNA-3). Although RNA-4 was not enriched for a BP, it is significantly associated with the cellular component ontology of histone deacetylase complexes, suggesting its role in early transcriptional regulation.

Together, these findings highlight time- and drug-dependent cardiomyocyte responses to TOP2i drugs at the level of individual genes and proteins and co-expressed modules. Therefore, this framework allows for an unbiased system level contextualization of known genes implicated in TOP2i cardiomyocyte dysfunction, as well as the exploration of less reported aspects of the molecular phenotype associated with CTRCD.

### Protein co-expression modules are preserved at the RNA level

By generating independent protein and RNA co-expression networks and assessing module eigenvalue correlations with TOP2i treatments, we identified time- and drug-dependent effects across molecular phenotypes. To determine if the co-expression relationships of drug-correlated modules in RNA translate to co-abundance relationships in protein, we tested for module preservation between networks. Specifically, we tested for the preservation of RNA modules in the protein network, and for the preservation of protein modules in the RNA network.

To assess module preservation, we utilized WGCNA’s Z-summary statistic to quantify similarities in the correlation structure between networks where modules with a Z-summary score > 2 are considered preserved. Preserved modules are further denoted as highly preserved (Z-score > 10) or moderately preserved (2 < Z-score < 10) based on the level of conservation of the RNA-protein relationships (Langfelder et al. 2011; Johnson et al. 2022). In the protein network we find that two thirds of co-expressed protein modules (12 of 18), and 80% of drug-correlated modules (8 of 10) are preserved in the RNA network (Fig 2A). Two modules (Pro-1 and Pro-2) are highly preserved in the RNA network and correspond to SDR modules 24 hours post drug treatment. In the RNA network we find that over half of co-expressed RNA modules (10 of 18) and two thirds of drug-correlated modules (7 of 11) are preserved in the protein network (Fig 2B). One module (RNA-2) is highly preserved in the protein network and corresponds to an SDR module at 24 hours. Given the tendency for drug-correlated modules to be preserved across networks, we asked whether there is a difference globally in the degree of preservation across SDR, VDR and NDR modules identified 24 hours post treatment. We observed that SDR modules have a higher level of preservation across networks than NDR modules (Wilcoxon-rank sum test*; P* < 0.05; Fig S9).

**Figure 2:**
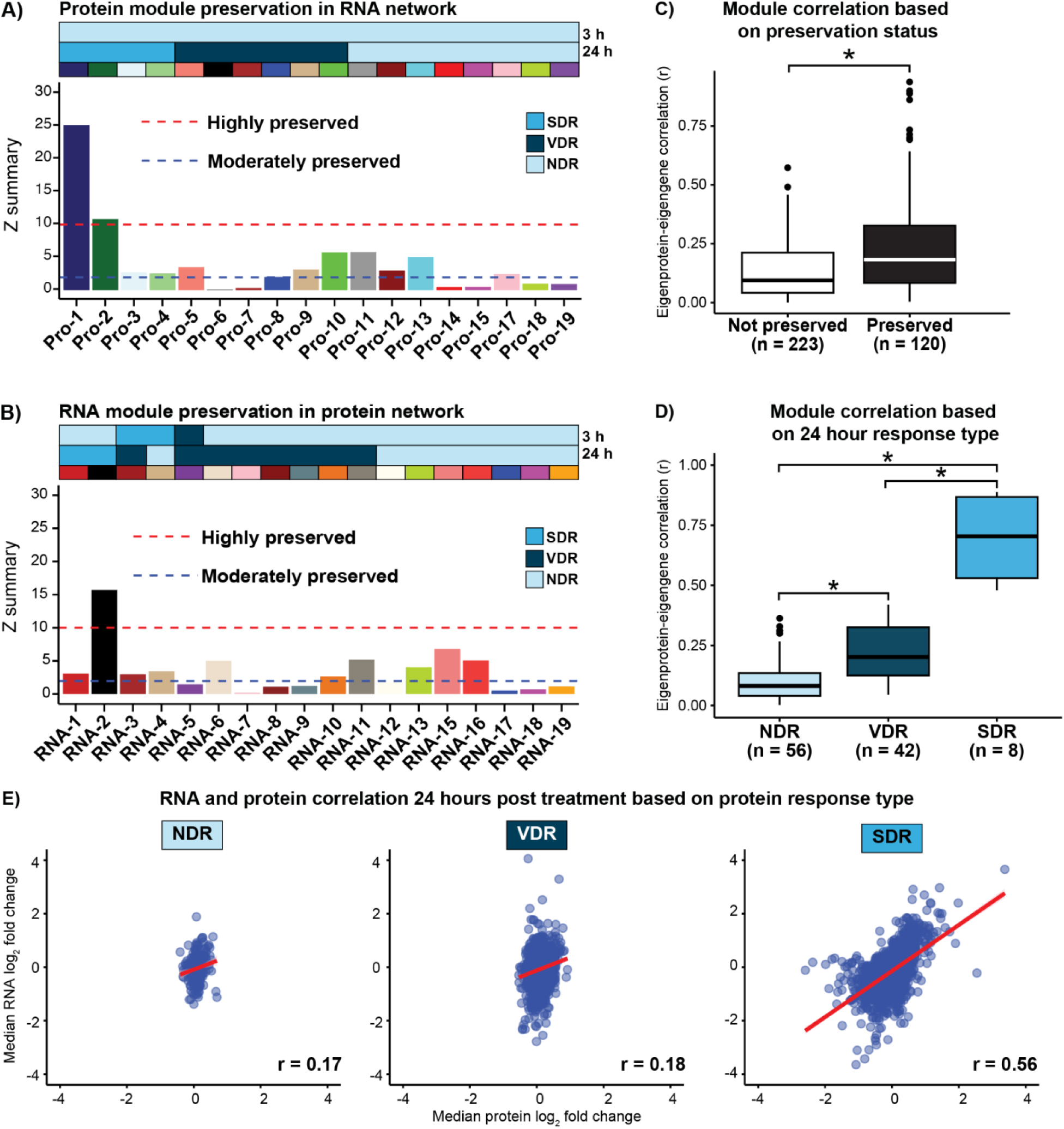
SDR protein modules are transcriptionally preserved. **A)** Module preservation results for each module in the protein network when testing for preservation of protein modules in the RNA network. Z summary statistics > 10 (red line) indicates strong preservation. Z summary statistics between 2 and 10 indicate moderate preservation (blue line). Z summary statistics < 2 indicate no evidence of preservation. Modules Pro-16 and RNA-14 are omitted, as these modules contain unassigned proteins/genes that are not co-expressed in any module, and therefore not given a Z summary statistic. **B)** Module preservation results for each module in the RNA network. **C)** Module eigengene-eigenprotein correlations based on preservation status (\**P* < 0.05). **D)** Comparison of eigengene-eigenprotein correlations across NDR, VDR and SDR modules (\**P* < 0.05). **E)** Pearson correlation between median drug responses between the overlapping set of proteins and their encoding genes in the RNA network at 24 hours post treatment in the NDR, VDR and SDR modules (\**P* < 0.05). Protein-gene pairs are assigned to modules based on their drug response in the protein data.

To directly compare modules in the protein and RNA networks, we calculated the Pearson correlation between every protein network eigenprotein and every RNA network eigengene and designated each module pair as being preserved if both the RNA and protein modules have Z-summary preservation scores > 2 and not preserved if neither RNA or protein module is considered preserved. We excluded unassigned modules from both networks (Pro-16 & RNA-14) since these modules do not represent co-expression relationships and therefore do not have a calculated Z-score. Preserved module pairs exhibited stronger eigengene-eigenprotein correlations compared to non-preserved module pairs (median r preserved = 0.18 and median r non-preserved = 0.09; Wilcoxon-rank sum test; *P* < 0.05; Fig 2C).

We next asked whether modules with similar responses to drug treatment in the RNA and protein networks also exhibit similar overall co-expression or co-abundance patterns. Specifically, we compared eigengene-eigenprotein correlations between SDR, VDR and NDR modules. We focused on the 24-hour module categorization given that it is the only timepoint where drug treatment is correlated with protein modules. We found that the 24-hour SDR modules show higher eigengene-eigenprotein correlations (median r = 0.70) compared to VDR modules (median r = 0.20) or NDR modules (median r = 0.08) indicating a greater similarity between RNA and protein responses in SDR modules (Wilcoxon-rank sum test; *P* < 0.05; Fig 2D). Similarly, VDR module eigengene-protein correlations are stronger than those of NDR modules (*P* < 0.05), suggesting that modules that respond to at least one drug maintain greater overall coordination between RNA and protein than those with no significant drug association.

We next determined whether the eigengene-eigenprotein correlation in drug-associated protein modules is reflected in the magnitude of the drug response in protein and RNA independent of the network structure. We calculated the correlation between the median log_2_ fold change in protein and RNA in response to drugs at the 24-hour timepoint in genes and proteins corresponding to the SDR, VDR and NDR protein modules. SDR modules show a strong log_2_ fold change correlation between protein and RNA (Pearson; r = 0.56; *P* < 0.05; Fig 2F). The protein-RNA response correlation is much lower in protein modules with lower drug association (VDR r = 0.18 and NDR r = 0.17; *P* < 0.05).

Together, these results suggest a coordinated response between RNA and protein in modules associated with all 24-hour drug treatments in comparison to modules associated with specific drug responses. This suggests that the shared molecular signatures across all drug treatments in the RNA and protein networks may be influenced by transcriptional control.

### Regulatory regions associated with SDR modules are enriched for stress-responsive transcription factor motifs

The high preservation of 24-hour SDR modules across protein and RNA networks suggests that transcriptional regulation likely drives the cellular response to drug treatment. We therefore sought to determine whether transcription factors (TFs) contribute to the regulation of genes in SDR modules. To identify candidate TFs, we obtained cleavage under targets & tagmentation (CUT&Tag) sequencing data for the H3K27ac histone modification, that marks active enhancers and promoters, in matched TOP2i-treated iPSC-CMs (Matthews et al. 2025).

We mapped 20,137 H3K27ac-enriched regions to the nearest gene transcription start site (TSS), and retained those associated with genes that are expressed in the RNA network (n = 17,163). To focus on the promoter regions of expressed genes, we selected H3K27ac-enriched regions within a 4 kb window around the TSS. This yielded 5,515 promoter-associated regions for downstream analysis. We tested for TF motif enrichment in RNA-1 and RNA-2 SDR module promoter-associated H3K27ac regions relative to H3K27ac-marked promoters from NDR module genes (Fig 3A; See Methods). TF enrichment analysis identified 146 significantly enriched TF motifs (Fisher’s exact test; 73log(adj. *P*) > 4; Table S20) associated with 133 TFs in RNA-2, and no enriched motifs in RNA-1 (Fig 3B). The most enriched TF motif corresponds to the basic helix-loop-helix protein BHLHE40 TF, and the motif for the canonical DNA damage response factor TP53 is amongst the most enriched motifs. Consistent with the association of H3K27ac with gene activation, RNA-2 is largely composed of DEGs that are upregulated in response to TOP2i, unlike RNA-1, which consists mostly of DEGs that are downregulated. RNA-2 is also highly preserved in the protein network supporting the role of these TFs in propagating molecular effects across RNA and protein.

**Figure 3:**
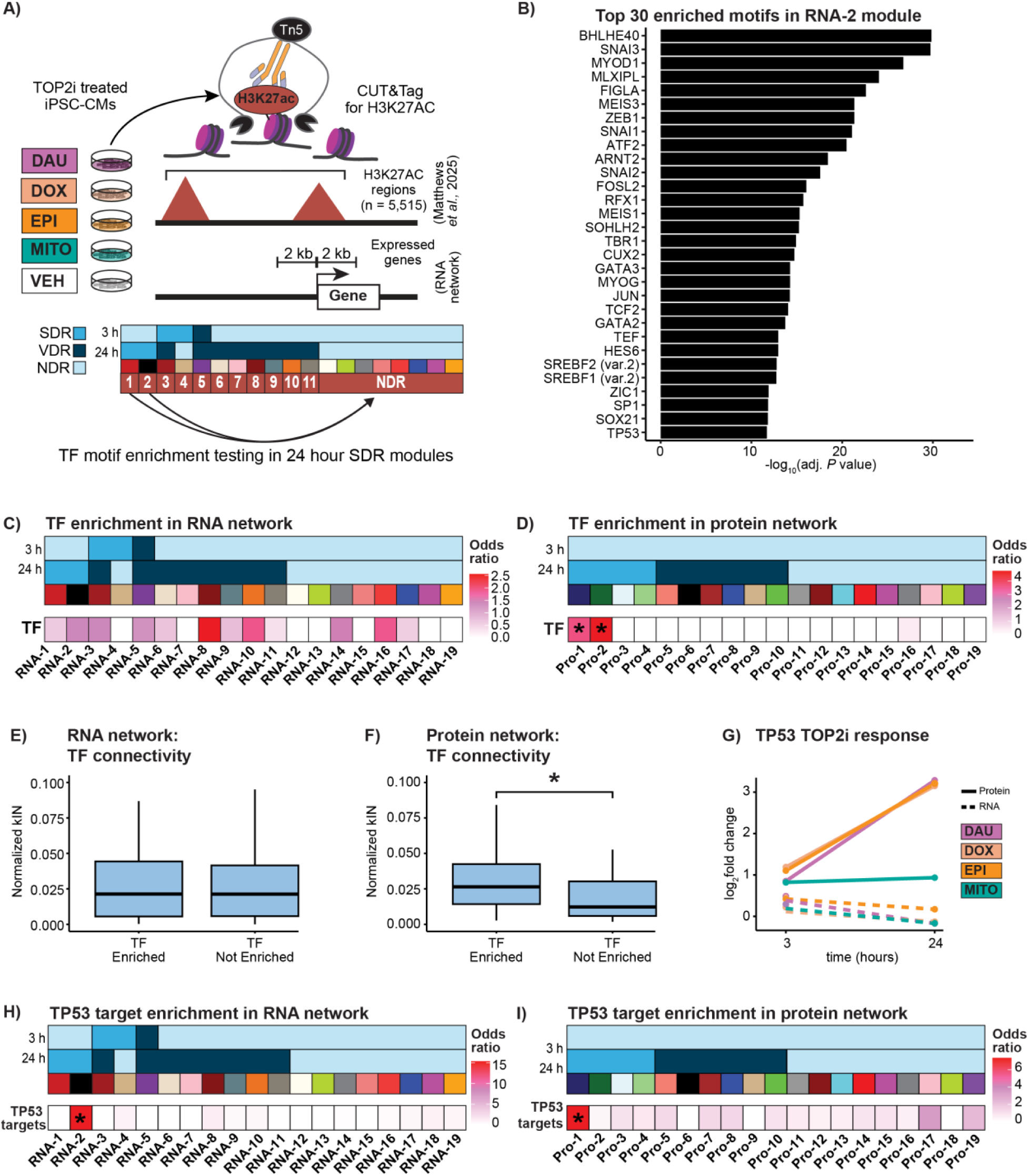
Preserved upregulated SDR module associates with the TP53 transcriptional regulator. **A)** Schematic illustrating the integration of H3K27ac CUT&Tag data from drug-treated iPSC-CMs (Matthews et al. 2025) and the RNA co-expression network to identify potential transcriptional regulators in the 24-hour SDR modules. Module RNA-2 is the only module enriched for transcription factor (TF) binding motifs. **B)** Top 30 enriched TF binding motifs (adj. *P* < 0.05) in the RNA-2 module. **C)** Enrichment of TFs that recognize the enriched TF motifs in modules of the RNA network (\**P* < 0.05). **D)** Enrichment of TFs that recognize the enriched TF motifs in modules of the protein network (\**P* < 0.05). **E)** Intra-modular connectivity (kIN) for TF genes associated with enriched TF binding motifs in the RNA network compared to TFs that are not associated with enriched motifs. **F)** kIN for TF proteins associated with enriched TF binding motifs in the protein network compared to TF proteins that are not associated with enriched motifs. **G)** TP53 response to TOP2i (log_2_ fold change compared to VEH) at the RNA level (dashed line) and protein level (solid line) over time. **H)** Enrichment of TP53 target genes within modules of the RNA network (\**P* < 0.05). **I)** Enrichment of TP53 target genes within modules of the protein network (\**P* < 0.05).

The majority of TFs that recognize the enriched TF motifs are expressed in the RNA network (61%; 82/133) providing further support for their involvement in the transcriptional response. We next asked if the TFs associated with enriched binding motifs in RNA-2 are co-expressed in the same module. We therefore tested for enrichment of the 82 expressed TF genes associated with enriched TF binding motifs in the RNA-2 module, across all modules in the RNA network. We find that TF genes associated with enriched TF binding motifs in the RNA-2 module are not enriched in any module in the RNA network (Fig 3C). We next tested for the enrichment of TF genes associated with enriched TF binding motifs in the RNA-2 module within the protein network. 17 of the 82 RNA expressed TFs associated with enriched motifs are also detected in the protein network. These TF proteins are enriched in the Pro-1 (Fisher’s-exact test; OR = 3.58; *P* < 0.05; n = 5) and Pro-2 (OR = 4.76; *P* < 0.05; n = 10) SDR modules, which are the only two protein modules to show high module preservation (Fig 3D). These data suggest a transcriptional network within RNA-2, Pro-1 and Pro-2 where TFs are co-expressed with genes associated with their binding motifs.

To determine how central TFs associated with the H3K27ac promoter regions are in the network, we assessed the connectivity of these TFs within the RNA and protein networks. Enriched TFs in the RNA network do not differ in their normalized intramodular connectivity (kIN) in comparison to non-enriched TFs (Fig 3E). However, TFs expressed at the protein level that are associated with enriched TF binding motifs have increased normalized kIN in the protein network compared to non-enriched TFs (Wilcoxon rank-sum test; *P* < 0.05; Fig 3F), suggesting that the TFs in the protein network are more central to their respective modules. This may in part be due to the differing dynamics of TF RNA and protein expression. For example, TP53 is a hub protein in the Pro-1 module and shows differential abundance in response to all drug treatments but is not differentially expressed at the RNA level (Fig 3G).

Given the central role of TP53 as a transcriptional regulator in our drug-treated network, we asked whether known TP53 target genes are enriched in drug-correlated modules (Fischer 2017). We find that genes activated by TP53 are enriched specifically in the RNA-2 module that is enriched for the TP53 motif (OR = 15.34; Chi-square test; *P* < 0.05; Fig 3H). TP53 targets are also specifically enriched in the Pro-1 protein module in which TP53 is a hub protein (OR = 7.36; *P* < 0.05; Fig 3I). These two modules have the highest module preservation between RNA and protein networks. These findings suggest that TF protein connectivity may serve as a stronger indicator of regulatory influence than transcriptional co-expression alone.

Collectively, these results indicate that TFs that associate with motifs enriched in H3K27ac-enriched regions are central to the shared drug response. These interactions indicate a highly preserved RNA-protein axis between modules RNA-2, Pro-1 and Pro-2 that is mediated by the TP53 hub protein.

### SDR protein modules link electrophysiological traits to impaired myocardial strain

Genome-wide association studies (GWAS) have identified variants associated with CVD risk, where genes in these loci are implicated in disease. However, whether these CVD risk genes influence the development of CTRCD is unclear. We therefore integrated proteins from drug-responsive modules with genes localized within CVD risk loci.

We first assessed whether modules in the protein network are enriched for genes mapped to loci for CVD or cardiovascular function (CF) related traits. We compiled 1,077 cardiovascular-related traits from the GWAS catalog including 993 related to CF measurements and 84 associated with CVDs and extracted the set of mapped genes (Cerezo et al. 2025). Two cardiovascular-related traits are enriched in our network (Chi-square test; adj. *P* < 0.05). The Pro-2 module is enriched for CVD risk proteins linked to atrial fibrillation (OR = 2.01; Fig 4A). This same module is enriched for CF proteins associated with PR-interval (OR = 2.35; Fig 4A). These two electrophysiological traits are closely related as PR interval dysregulation can contribute to atrial fibrillation, which is characterized by the absence of discernible P waves on an electrocardiogram (Cheng et al. 2009; Cheng et al. 2015).

**Figure 4:**
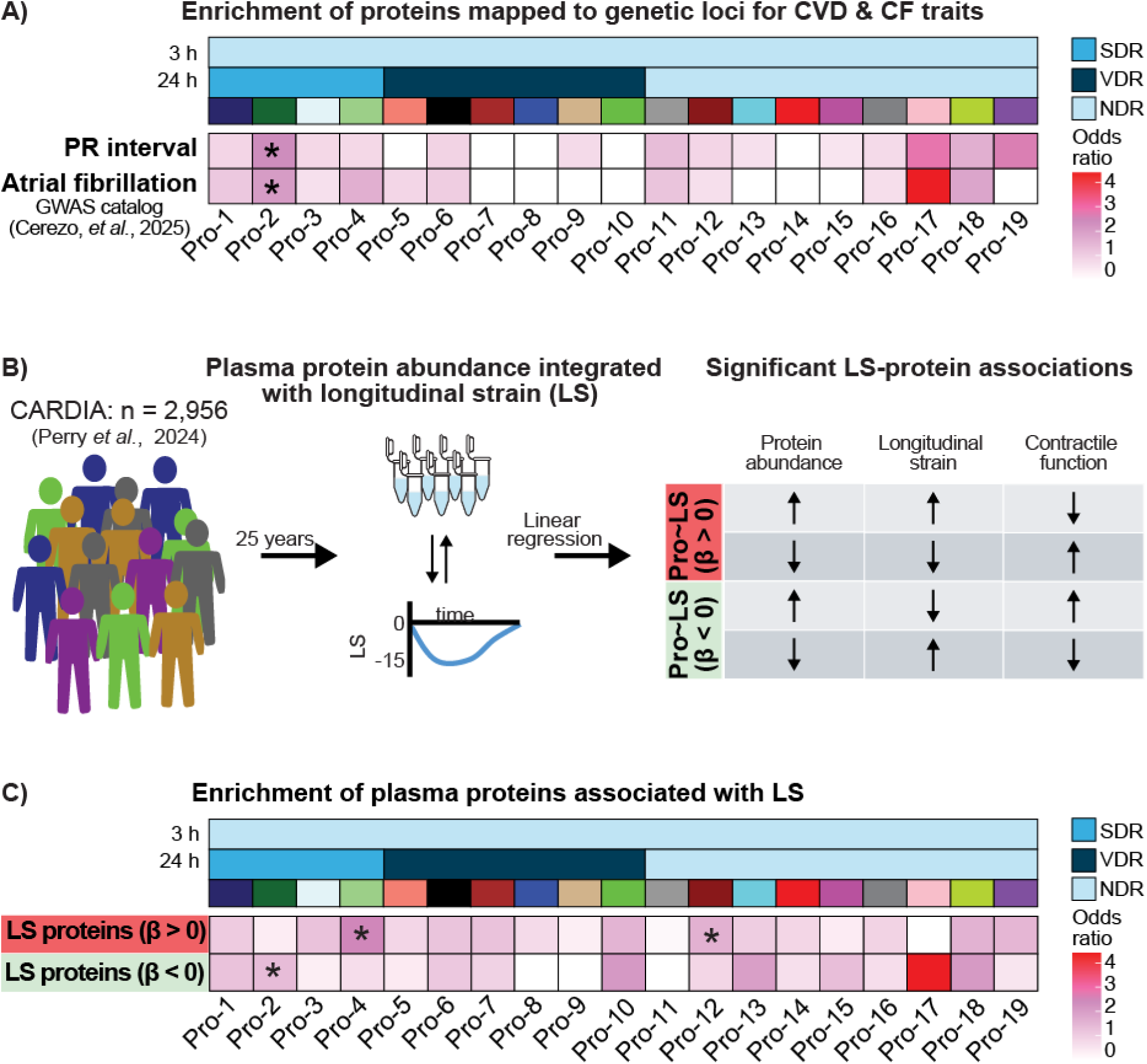
SDR protein modules associate with arrhythmogenic and CVD traits, and proteins that influence myocardial strain. **A)** Enrichment of genes in loci associated with CVDs and cardiac function (CF) within modules of the protein network (*adj. *P* < 0.05) (Cerezo et al. 2025). **B)** Schematic showing the identification of proteins that associate with longitudinal strain (LS) from the CARDIA study (Perry et al. 2024). Potential directions of the LS-protein association are illustrated together with the inferred effects on contractility. **C)** Enrichment of genes associated with proteins with a significant positive association with LS and negative association with LS within modules of the protein network (*adj. *P* < 0.05).

We next examined specific genes associated with atrial fibrillation and PR interval that are contained in the Pro-2 module. NKX2-5, a known atrial fibrillation risk gene (Nielsen et al. 2018; Assum et al. 2022), and MEIS1, associated with PR interval (Pfeufer et al. 2010), are both TFs whose motifs are enriched in H3K27ac-enriched promoter sequences in the RNA-2 module. Their protein expression is decreased in response to all AC drug treatments (Fig S10). The Pro-2 module also contains seven hub proteins associated with atrial fibrillation risk (GTF2I, CGNL1, LRRC10, NCOR2, SLIT3, AKAP6, and CASZ1) (Roselli et al. 2018; Roselli et al. 2025) and seven hub proteins linked to PR interval (NGDN, SIPA1L1, CBX8, AKAP6, ADGRL3, SIPA1L2, and FNDC3B) (Van Setten et al. 2018; Ntalla et al. 2020). These results suggest that the Pro-2 module may serve as an important regulatory module integrating TF activity, contractility traits, and CVD risk gene networks in the context of TOP2i-induced cardiotoxicity.

To further investigate how the response to TOP2i treatment can influence cardiomyocyte contractility, we integrated our protein network data with proteins associated with a clinical metric used for early detection of CTRCD - longitudinal strain (LS) (Thavendiranathan et al. 2014; Liu et al. 2020). LS reflects the ability of the myocardium to contract and shorten along its longitudinal axis and provides a more nuanced assessment of systolic function than traditional ejection fraction measurements. Identifying treatment-correlated protein modules enriched for LS-associated proteins can therefore provide insights into the molecular effects coupled to early myocardial contractile dysfunction in CTRCD.

To assess the enrichment of LS-associated proteins within modules of the protein network, we obtained LS-associated proteins identified by Perry *et al*. (Perry et al. 2024). Highly negative LS scores indicate normal cardiac contractile function while less negative scores indicate cardiac contractile dysfunction. Perry and colleagues analyzed proteomics data from plasma samples and LS from 2,956 individuals in the Coronary Artery Risk Development in Young Adults (CARDIA) cohort (Friedman et al. 1988; Lloyd-Jones et al. 2021). The data analyzed was obtained from participants who attended a follow-up visit 25 years after baseline protein and LS measurements were gathered. These individuals (mean age = 50 years at follow-up; 56% women) show an overall low prevalence of CVD and moderate to high prevalence of risk factors across the cohort. By regressing protein expression levels against LS measurements they identified proteins with significant positive (β > 0) and negative (β < 0) associations with myocardial strain. We integrated these data with our TOP2i-treated iPSC-CM protein co-abundance network (Fig 4B). We find that the Pro-2 module is enriched for proteins negatively associated with LS (OR = 1.48; Chi-square test; adj. *P* < 0.05; Fig 4C). This is a set of proteins where an increase in their expression associates with reduced LS score and better cardiac contractile function. Pro-2 contains proteins exhibiting decreased abundance in response to all drug treatments at 24 hours, consistent with an adverse effect on cardiac contractile function. Conversely, Pro-4 (OR = 2.70; adj. *P* < 0.05) and Pro-12 (OR = 1.71; adj. *P* < 0.05;) are enriched for proteins positively associated with LS (Fig 4C). This is a set of proteins where an increase in their expression associates with an increased LS score, leading to worse cardiac contractile function. Pro-4, which is positively correlated with all drug treatments at 24 hours, contains proteins that show increased abundance due to treatment, again consistent with an adverse effect on cardiac contractile function. These results suggest that 24-hour SDR protein modules Pro-2 and Pro-4 contain proteins linked to impaired myocardial contractility, and parse LS-associated proteins responding to TOP2i treatment from those that do not.

These findings reveal that TOP2i-treated cardiomyocytes exhibit molecular signatures linked to arrhythmogenic traits and impaired myocardial strain. Specifically, the Pro-2 module emerges as a module enriched for both atrial fibrillation and PR interval risk proteins that act as TFs, as well as proteins negatively associated with LS, suggesting a role in chemotherapy-induced electrophysiological and contractile dysfunction.

### Preserved downregulated SDR protein module is linked to cytosolic calcium dynamics

Because arrhythmogenic risk genes and myocardial strain–associated proteins are enriched in the SDR module, Pro-2, we hypothesized that these proteins might affect the contractile function of iPSC-CMs. Cardiomyocyte contraction is tightly coupled to cytosolic calcium dynamics. We therefore obtained calcium transient data from iPSC-CMs from three of the individuals used in this study that were treated with DAU, DOX, EPI, MITO or VEH for 24 hours (Matthews et al. 2024). We collated data on the rate at which calcium transients occur, as well as features of the transients such as their amplitude, rise slope (10%-90% amplitude), and decay slope (90%-10% amplitude; Fig 5A). All of these calcium transient features are affected by at least one TOP2i treatment (t-test; *P* < 0.05; Fig 5B). The decay slope is significantly prolonged compared to VEH across all drug treatments suggesting a decreased rate of calcium efflux from the cytosol.

**Figure 5:**
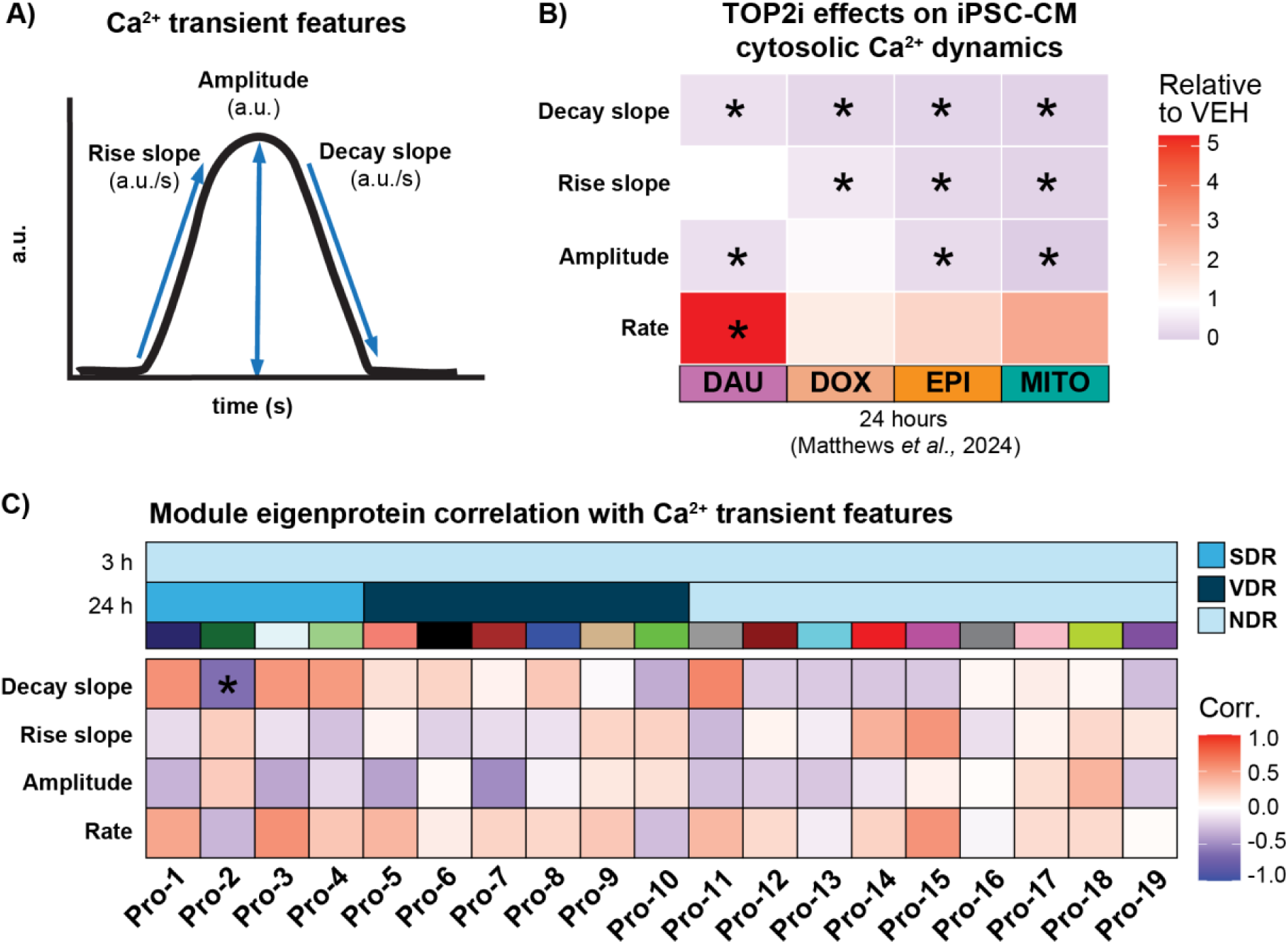
Pro-2 SDR module is correlated with cytosolic calcium efflux. **A)** Schematic representation of the features obtained to assess intracellular calcium flux in beating iPSC-CMs from three individuals treated with DAU, DOX, EPI, MITO and VEH for 24 hours (Matthews et al. 2024). Fluorescent intensity of the calcium-sensitive fluorescent dye Fluo-4 AM was measured over time by spinning disc confocal microscopy. **B)** Mean calcium transient amplitude, rise slope, decay slope and rate for every treatment at 24 hours relative to VEH (\**P* < 0.05). Shading determined by median value in each drug treatment normalized to the VEH median. **C)** Pearson correlation of module eigenproteins to standardized values for mean amplitude, rise slope, decay slope and rate across TOP2i-treated samples (*adj. *P* < 0.05).

We next asked whether the 24-hour drug–induced changes in calcium transients align with specific protein network modules. Because cytosolic calcium flux is ultimately controlled by ion-handling proteins, we used module eigenproteins rather than eigengenes to link molecular signatures to calcium dynamics. For every module we kept only those eigenprotein values corresponding to samples in which calcium-transient features (rise slope, decay slope, amplitude) were recorded. Pearson correlations were independently calculated between standardized eigenprotein values and each standardized calcium-transient metric for each module. After multiple testing adjustment, a significant correlation indicates that the proteins grouped in that module track with the functional calcium response to drug treatment after 24 hours of treatment. The SDR module Pro-2 is the only module that is correlated with cytosolic calcium handling, showing a negative correlation with the calcium transient decay slope (r = −0.63; adj. *P* < 0.05; Fig 5C). This module contains proteins that are downregulated in response to drug treatment. Because the decay slope becomes less negative across all treatments, indicating a slower rate of cytosolic calcium efflux, the negative correlation between Pro-2 and the decay slope suggests that decreased protein abundance in this module is associated with impaired calcium clearance.

Together, these findings link molecular responses to drug treatment in the Pro-2 SDR protein module with functional disruptions in cardiomyocyte calcium handling, thereby providing a molecular context for early CTRCD.

## DISCUSSION

Despite the well-established cardiotoxic effects of TOP2i there remains a critical gap in understanding the early molecular events that precede overt cardiomyocyte death, and connecting these effects with risk for developing cardiovascular disease (Joglar et al. 2024). In this study, we established an iPSC-CM model across six individuals to investigate the early molecular disruptions induced by TOP2i chemotherapeutics prior to cell death using sub-micromolar, clinically relevant concentrations of DAU, DOX, EPI and MITO. We identified drug-responsive co-expression modules that are preserved across RNA and protein networks, link to gene regulatory factors, and associate with PR interval and atrial fibrillation risk.

Construction of RNA and protein co-expression networks in cardiomyocytes treated with anthracyclines revealed both time- and drug-dependent molecular signatures that emerge following treatment with TOP2i. Signatures characterized by treatment responses across all drugs, SDR modules, are more likely to be preserved across RNA and protein networks compared to VDR modules that show variability in the response to treatment across drugs. This divergence suggests that SDR protein modules are governed by transcriptional regulation, while VDR protein modules may be more strongly influenced by post-transcriptional, post-translational or other protein-level regulation.

By using data identifying H3K27ac-enriched regions in response to TOP2i (Matthews et al. 2025), we identified TF motifs enriched in SDR RNA module promoters, which correspond to TFs expressed in networks that have been reported to influence CTRCD, including TP53, GATA4, NKX2-5, SP1 and STAT1 (Aries et al. 2004; Nakamura et al. 2019; Saleme et al. 2019; Sada et al. 2023; Xiang et al. 2025). We also find motifs for treatment-responsive TFs that have not been associated with cardiotoxicity, such as hub protein SREBF2, and DAP TFs such as MEIS1 and JUNB. Several of these TFs, including the hub protein TP53, exhibit distinct treatment response dynamics at the RNA and protein levels, suggesting a role for post-transcriptional control that may not be captured at the level of RNA expression alone. While TP53 expression and abundance exhibit distinct treatment responses, TP53 protein is co-abundant with its predicted gene targets. These data reinforce a regulatory axis that emerges when integrating information across transcriptional activation, gene expression and protein expression in response to TOP2i treatment.

In the protein network, SDR module Pro-2 contains proteins with decreased abundance due to treatment at 24 hours and are most enriched for biological processes related to transcriptional regulation, RNA processing, and ribosome biogenesis. The only three DAPs across all TOP2i and timepoints are POLR2A, FCF1, and hub protein TBL3 – all of which participate in transcriptional regulation, RNA processing, and ribosome biogenesis. However, the response to treatment differs at the level of RNA. POLR2A is localized within SDR module RNA-1 and is an upregulated DEG in all 24-hour treatments, but a downregulated DAP in all treatments. FCF1 is a DEG only after 24 hours of treatment with EPI, and TBL3 is not differentially expressed as RNA under any condition. Taken together these observations show that Pro-2 is markedly enriched for proteins related to ribosome-biogenesis, suggesting that disrupted ribosomal assembly may be a contributor to early cardiomyocyte injury and dysfunction. These results are in line with the observation that nucleolar stress is associated with early myocardial damage (Avitabile et al. 2011).

The Pro-2 module, a highly preserved SDR module, is significantly enriched for genes linked to genetic risk for atrial fibrillation and PR interval, two closely related electrophysiological traits (Joglar et al. 2024). These data suggested the potential for contractile dysfunction. However, atrial fibrillation and PR interval are traits that correspond to the coordination of multiple cell types and are not limited to cardiomyocytes alone (Joglar et al. 2024). We therefore tested for association of module proteins with those that influence LS, a feature of myofibrillar contraction (Thavendiranathan et al. 2014; Liu et al. 2020). We found that Pro-2 and Pro-4 are enriched for LS-associated proteins with a direction of effect at 24 hours indicative of disrupted myocardial function (Perry et al. 2024). Functional assessment of cardiomyocyte contractility (Matthews et al. 2024) identified that calcium efflux from the cytosol was slowed across all TOP2i treatments at 24 hours, indicating functional impairment to contraction. Differences in calcium efflux correlate with the eigenprotein in Pro-2. Proteins with known roles in cardiomyocyte contraction regulation reside in this module, including sodium voltage-gated channel protein type 5 (SCN5A), which is downregulated following treatment with all TOP2i treatments at 24 hours.

The high preservation of perturbed RNA and protein networks in cardiomyocytes and their link to complex cardiovascular disease risk genes that we observed should be considered in the context of multi-omic studies in other disease systems. Paired RNA and protein networks from Alzheimer’s and healthy brain tissue have indicated that the association with Alzheimer’s’ disease is linked to modules uniquely identified in the protein network (Seyfried et al. 2017). This difference in the contribution of mRNA changes to disease pathology is likely indicative of disease-specific etiologies. However, genes in regions in loci associated with Alzheimer’s disease often localized in modules present in the RNA and protein networks suggesting mechanistic pathways of disease association. Similarly, genes implicated in genetic risk for Alzheimer’s disease are enriched in protein co-expression modules that are associated with early onset dementia in line with relevant GWAS genes being enriched in our early DNA damage response modules (Swarup et al. 2020).

Clinically, cardiotoxicity detection relies largely on cardiac troponin T and troponin I release, which are elevated after cardiomyocyte injury has already occurred. Alternatively, LS measurements offer early functional readouts that are highly sensitive and specific to one’s CTRCD risk prior to cardiomyocyte death. However, the underlying molecular mechanisms behind CTRCD-associated contractile impairments, rather than cell death, have been minimally characterized. Given our protein network’s association with LS, atrial fibrillation and disrupted calcium handling at sub-lethal TOP2i concentrations, SDR protein modules may be used to further investigate biomarkers indicative of early risk of contractile dysfunction due to TOP2i treatment.

Similarly, a pipeline for discovering therapies that can prevent cardiotoxicity before irreversible injury occurs is still needed. Molecular studies can identify modifiers of DOX toxicity. For example, a CRISPR-i/a screen of thousands of genes in iPSC-CMs by Liu *et al*. highlighted the role of CA12 (Liu et al. 2024). Notably this gene is a hub gene in our SDR module RNA-2. However, the screen was restricted to genes already considered to be druggable and evaluated only a single AC. Our multi-omic analysis offers a broader and less biased vantage point to identify specific gene targets and pathways for further mechanistic follow-up. The SDR modules we identified are preserved between RNA and protein levels and are activated by every TOP2i we tested, implying that modulating SDR hubs, shared across multiple drugs at the RNA level may be more likely to propagate to protein. By focusing on the response of all expressed iPSC-CM genes and detectable proteins to drug treatment, we provide data to move beyond the currently limited “druggable genome” and toward transcript-level interventions that impact protein levels and pre-empt cardiotoxicity across multiple chemotherapeutic drugs.

While this study provides a detailed multi-omic framework to understand early molecular events in CTRCD, several limitations should be considered. First, although mass spectrometry is a powerful tool for quantifying protein abundance, low-abundance proteins may be underrepresented and lead to an incomplete protein abundance profile. Second, while we collected RNA and protein from the same differentiation and treatment batch, different cells were interrogated for each molecular phenotype. Single cell multi-omic studies would allow for more direct comparison of gene regulatory layers (Leduc et al. 2025). Third, we investigated the direct cardiomyocyte response to drugs; however the response may be modulated by surrounding cell types including endothelial cells, fibroblasts and immune cells.

In summary, we integrated global mRNA and protein expression data and epigenetic profiling from human cardiomyocytes treated with TOP2i with genetically-associated cardiovascular traits and direct measures of contractile function. We identified a drug-responsive protein co-expression module that is transcriptionally preserved, and is associated with atrial fibrillation risk and myocardial dysfunction. Collectively, this systems-level dataset provides a resource for future studies seeking to identify early drug-induced cardiotoxicity biomarkers and therapeutic targets to prevent irreversible myocardial damage.

## METHODS

### Induced pluripotent stem cell lines

iPSC lines were selected from the iPSCORE resource that was generated by Dr. Kelly A. Frazer at the University of California San Diego as part of the National Heart, Lung and Blood Institute Next Generation Consortium (Panopoulos et al. 2017). The iPSC lines were generated with approval from the Institutional Review Boards of the University of California, San Diego and The Salk Institute (Project no. 110776ZF) and informed written consent of participants. Cell lines are available through the biorepository at WiCell Research Institute (Madison, WI, USA), or through contacting Dr. Kelly A. Frazer at the University of California, San Diego.

We used iPSC lines from six unrelated, healthy female donors of Asian ethnicity between the ages of 21 and 32 years with no previous history of cardiac disease or cancer. The individuals are: Individual 1: UCSD129i-75-1 (iPSCORE_75_1, Asian-Irani, age 30), Individual 2: UCSD143i-87-1 (iPSCORE_87_1, Asian-Chinese, age 21), Individual 3: UCSD131i-77-1 (iPSCORE_77_1, Asian-Chinese, age 23), Individual 4: UCSD133i-79-1 (iPSCORE_79_1, Asian, age 24), Individual 5: UCSD132i-78-1 (iPSCORE_78_1, Asian-Chinese, age 21), and Individual 6: UCSD116i-71-1 (iPSCORE_71_1, Asian, age 32).

### Generation of proteomics data

#### iPSC culture

Feeder-free iPSCs were cultured at 37 °C, 5% CO_2_ and atmospheric O_2_ in mTESR1 (85850, Stem Cell Technology, Vancouver, BC, Canada). The mTESR1 media included 1% Penicillin/Streptomycin (30-002-Cl, Corning, Bedford, MA. USA). Cells were cultured on a substrate containing a 1:100 dilution of Matrigel hESC-qualified Matrix (354277, Corning). Sub-confluent iPSCs were passaged using dissociation reagent (0.5 mM EDTA, 300 mM NaCl in PBS) every 3–5 days.

#### Cardiomyocyte differentiation from iPSCs

Cardiomyocyte differentiation was performed as previously described (Matthews et al. 2024). Briefly, Cardiomyocyte Differentiation Media (CDM) containing 12 μM CHIR99021 trihydrochloride (4953, Tocris Bioscience, Bristol, UK) was added to 80-95% confluent iPSCs on Day 0. Twenty-four hours later, the media was replaced with CDM. After 48 hours, media was replaced with CDM containing 2 μM Wnt-C59 (5148, Tocris Bioscience). CDM was added on Days 5, 7, 10, and 12. Cardiomyocytes were purified through metabolic selection on Days 14,16 and 18 using glucose-free Purification Media. On Day 20, 1.5 million iPSC-CMs were replated per well of a six-well plate, in glucose-free Cardiomyocyte Maintenance Media (CMM). iPSC-CMs were matured in culture with CMM replaced on Days 23, 25, 27, 28, and 30.

#### Determination of iPSC-CM purity by flow cytometry

After each iPSC differentiation, live cardiomyocyte purity was assessed using flow cytometry as previously described (Matthews et al. 2024). Day 25–27, iPSC-CMs were stained with a live-dead stain (Zombie Violet Fixable Viability Kit (423113, BioLegend, San Diego, CA, USA) and cardiac troponin T antibody (Cardiac Troponin T Mouse, PE, Clone: 13–11, BD Mouse Monoclonal Antibody (564767, BD Biosciences, San Jose, CA, USA)). Control samples included unlabeled iPSC-CMs and iPSC-CMs stained for either Zombie or cardiac troponin T antibody only. Ten thousand cells were analyzed per sample on a BD LSR Fortessa Cell Analyzer. Data presented as the mean percentage of live, cardiac troponin T-positive cells across two technical replicates.

#### Drug treatment of iPSC-CMs

Between Days 27–29, iPSC-CMs were treated with 0.5 μM DAU (30450, Sigma-Aldrich, Saint Louis, MO, USA), DOX (D1515, Sigma-Aldrich), EPI (E9406, Sigma-Aldrich), MITO (M6545, Sigma-Aldrich), or a water vehicle (VEH) in CMM and collected three- and 24-hours post-treatment. iPSC-CMs were washed twice with ice-cold PBS, manually scraped in cold PBS on ice, and cell pellets flash-frozen and stored at −80 °C.

#### Cell viability determination

55,000 Day20 iPSC-CMs from Individual 5 were plated per well of a 96 well plate. On Day 27, cells were treated with 0.5 µM of DAU, DOX, EPI, MITO, or VEH control for 24 hours in quadruplicate. Sample positions on the plate were randomized using Well Plate Maker (wpm) package in R (Borges et al. 2021). 24 hours after treatment, media was aspirated, and cells were washed two times with DPBS. PrestoBlue Cell Viability reagent (A13261, Invitrogen, Thermo Fisher Scientific, San Jose, CA, USA) was added to treated cells according to manufacturer’s instructions. Plate reading was performed using a Biotek Synergy H1 (Agilent, Thermo Fisher Scientific) plate reader set to an excitation/emission of 560/590 nm. Data was processed using R according to manufacturer’s instructions. Briefly, background fluorescence measured from wells containing Cell Viability reagent only was averaged (n = 4) and subtracted from all sample wells, yielding a relative fluorescence unit (RFLU) value for each sample. Each drug-treated sample was divided by the average RFLU from all VEH-treated wells (n = 4) and multiplied by 100 to generate a percent cell viability for each sample. A t-test was used to determine differences between drug and VEH treatments.

#### Protein isolation and quantification

60 iPSC-CM samples were arranged into ten protein extraction batches corresponding to all treatment samples per individual. Protein was isolated by lysing the cells with 100 µl ice-cold RIPA buffer [1.5 ml 5 M NaCl, 1 ml Triton X100, 1 g Na deoxycholate, 1 ml 10% SDS, 1 ml 1 M Tris pH 7.4 and 45 ml dH_2_O] with protease inhibitor for 1 h at 4 °C. Post cell lysis the protein contents of the cells were separated from the lysed cell debris by centrifuging at 15,000 g for 15 min at 4°C. The protein-containing supernatant was stored at −80 °C. The isolated proteins were quantified in duplicate with the BCA Protein Assay kit (23227, Thermo Scientific) according to the manufacturer’s instructions

#### Protein digestion

We aimed to limit batch effects in the protein digestion process through block randomization (Burger et al. 2020). The 60 extracted protein samples were organized into three randomized blocks, where each block contained protein samples across all treatments from two individuals. Samples within each 20-sample block were randomized. Protein digestion was performed in these randomized batches. The samples were digested similarly as previously described (Herrmann et al. 2021). Briefly, 15 μg of protein were solubilized with 60 μL of 50 mM Triethylammonium bicarbonate (TEAB) pH 7.55. The proteins were then reduced with 10 mM Tris(2-carboxyethyl) phosphine (TCEP) (77720, Thermo) and incubated at 65 °C for 10 min. The sample was then cooled to room temperature and 1 μL of 500 mM iodoacetamide acid was added and allowed to react for 30 min in the dark. Then, 3.3 μl of 12% phosphoric acid was added to the protein solution followed by 200 μL of binding buffer (90% Methanol, 100mM TEAB pH 8.5). The resulting solution was added to S-Trap spin column (protifi.com) and passed through the column using a bench top centrifuge (60 s spin at 1,000 g). The spin column is washed with 150 μL of binding buffer and centrifuged. This is repeated two times. 30 μL of 20 ng/μL Trypsin is added to the protein mixture in 50 mM TEAB pH 8.5, and incubated at 37 ^○^C overnight. Peptides were eluted twice with 75 μL of 50% acetonitrile, 0.1% formic acid. Aliquots of 20 μL of eluted peptides were quantified using the Quantitative Fluorometric Peptide Assay (23290, Pierce, Thermo Fisher Scientific). Eluted volume of peptides corresponding to 5.5 μg of peptides are dried in a speed vac and resuspended in 27.5 μL 1.67% acetonitrile, 0.08% formic acid, 0.83% acetic acid, 97.42% water and placed in an autosampler vial.

#### Nanoflow liquid chromatography mass spectrometry

Peptide mixtures were analyzed by nanoflow liquid chromatography-tandem mass spectrometry (LC-MS/MS) using a nano-LC chromatography system (UltiMate 3000 RSLCnano, Dionex, Sunnyvale, CA, USA), coupled on-line to a Thermo Orbitrap Eclipse mass spectrometer (Thermo Fisher Scientific) through a nanospray ion source. Instrument performance was verified by analyzing a standard six protein mix digest before the sample set run, between each experimental block and at the end of the experiment. Protein batches were block randomized by individual, where batch contained sequentially randomized samples from each individual (n = 10). One protein mix digest of all samples in the individual block (10-pool sample) was run before and after all randomized samples. We further ran a protein mix digest comprising all 60 experimental samples at the end of each run (60-pool sample). The protein mix data files were analyzed to confirm that instrument performance remained consistent throughout the experiment. In total 60 experimental samples, 17 x10-sample batch pooled samples, 9 x 60-sample pooled samples, and 6 rerun experimental samples were processed. This led to a total of 92 processed samples. A direct injection method using 3 μL of digest onto an analytical column was used (Aurora (75 µm X 25 cm, 1.6 µm, IonOpticks, Collingwood, VIC, Australia). After equilibrating the column in 98% solvent A (0.1% formic acid in water) and 2% solvent B (0.1% formic acid in acetonitrile (ACN)), the samples (2 µL in solvent A) were injected (300 nL/min) by gradient elution onto the C18 column as follows: isocratic at 2% B, 0-5 min; 2% to 5%, 5-5.1 min; 5% to 19% 5.1-60 min, 19% to 35% B, 60-65 min; 50% to 90% B, 65-68 min; isocratic at 90% B, 68-70 min; 90% to 10%, 70-71 min; isocratic at 10% B, 71-72 min; 10% to 95% 72-73 min; isocratic for two min; 95%-2%, 73-76 min; and isocratic at 2% B, till 90 min.

#### Nanoflow liquid chromatography mass spectrometry analysis for DIA

All LC-MS/MS data were acquired using an Orbitrap Eclipse in positive ion mode using a data-independent acquisition (DIA) method with 8 Da windows from 400-900 and a loop time of 3 s. The survey scans (m/z 350-2000) were acquired in the Orbitrap at 60,000 resolution (at m/z = 400) in centroid mode, with a maximum injection time of 118 ms and an AGC target of 100,000 ions. The S-lens RF level was set to 60. Isolation was performed in the quadrupole, and HCD MS/MS acquisition was performed in profile mode using the orbitrap at a resolution of 30000 using the following settings: collision energy = 33%, IT 54 ms, AGC target = 50,000. A pooled sample was used to create spectral libraries that we search the individual samples against by injecting 5 times using narrow (4 Da), staggered windows over 100 m/z ranges from 400-900 m/z in a technique called gas phase fractionation as described in Searle *et al*. (Searle et al. 2020).

#### Database searching for DIA proteins

The raw data was demultiplexed to mzML with 10 ppm accuracy after peak picking in MSConvert (Chambers et al. 2012). The resulting mzML files were searched in MSFragger (Kong et al. 2017) and quantified via DIA-NN (Demichev et al. 2020) using the following settings: peptide length range 7-50, protease set to Trypsin, 1 missed cleavage, 3 variable modifications, clip N-term M on, fixed C carbamidomethylation, variable modifications of methionine oxidation and n-terminal acetylation, MS1 and MS2 accuracy set to 20 ppm, 1% FDR, and DIANN quantification strategy set to Robust LC (high accuracy). The files were searched against a database of human acquired from Uniprot (18^th^ December, 2023). The gas-phase fractions were used only to generate the spectral library, which was used for analysis of the individual samples.

#### Analysis of proteomics and RNA-seq data

All custom analysis scripts used in this project are available at https://mward-lab.github.io/Johnson_multiomic_network_2025/index.html made possible by workflowr (Blischak et al. 2019).

#### Determination of global mRNA expression levels

RNA-seq data was obtained from our previous work described in Matthews *et al*. (Matthews et al. 2024). Briefly, iPSC-CMs from the same differentiation experiments described for protein were drug-treated and collected at the same time for RNA extraction. RNA was extracted in six batches where all samples per individual were extracted together. The median RIN score across samples was 9.6. RNA-seq libraries were generated in six batches corresponding to all samples per individual. All samples were sequenced as a single pool. We used the RNA counts file (GEO: GSE243674) as the basis for the analysis described in this paper. We transformed the counts to log_2_ counts per million with the edgeR package and excluded genes with a mean log_2_ cpm < 0 across samples, leaving 14,128 expressed genes for downstream analysis (Chen et al. 2016).

#### Determination of global protein abundance levels

##### Abundance matrix filtering and outlier removal

Following LC-MS/MS we had protein abundance data for 60 experimental samples, 10-sample batch pools run at the beginning and end of the batch, and 60-sample pools run for each batch. When comparing within batch pools at the beginning and end of the LC-MS/MS run, we observed overlapping abundance distributions indicating an absence of machine drift at this timescale.

We identified 7,628 proteins present in at least one of the 60 experimental samples. We removed two non-human or uncharacterized proteins. Proteins that were missing in any one of the 60 experimental samples were removed, leaving 5,231 analyzable proteins. No imputation was performed regardless of type of missingness of proteins in the data. Hierarchical clustering of protein log_2_ abundances showed one outlier sample (Ind_1_DOX_3hrs), which was subsequently removed. These steps led to a total of 5,231 analyzable proteins across 59 samples to generate a log_2_-transformed abundance matrix for subsequent downstream analyses.

##### Removal of unwanted technical variation and normalization

To eliminate unwanted technical variation that may have arisen during the LC-MS/MS experiment, we combined our experimental sample data with the 60-sample pool data and adjusted the log_2_-transformed abundance matrix using both negative control proteins and the pooled protein samples comprised of all experimental samples that were included in every mass spectrometry batch i.e. technical replicates. We used the Remove Unwanted Variation (RUV) R package and RUV-III function where negative control proteins were defined as the 8% least variable proteins across all samples (Gagnon-Bartsch and Speed 2012). The 60-sample pool data was then removed, and only the 59 experimental samples remained. This analysis resulted in a RUV-corrected abundance matrix.

#### Protein co-abundance network generation using Weighted Gene Co-expression Network Analysis

We adopted the Weighted Gene Co-expression Network Analysis (WGCNA) methodology to investigate correlations between protein abundances in our RUV-corrected abundance matrix of DOX-, DAU-, EPI-, MITO- and VEH-treated iPSC-CMs collected three and 24 hours post-treatment (Langfelder and Horvath 2008). The WGCNA framework, primarily utilizing wrapper functions from BioNERO, was implemented for this analysis (Almeida-Silva and Venancio 2022). We first established a scale-free network topology, achieved by determining the appropriate soft threshold power using the SFT_fit function. After iterating different soft power thresholds (β), the linear regression of log_10_(k) versus log_10_(p(k)) indicates that by setting β = 10, the network is close to a scale-free network, where k is the whole network connectivity and p(k) is the corresponding frequency distribution. This setting resulted in our protein network attaining a scale-free fit index of 0.83 (quantification of how well the network approximates a scale-free topology), with a mean/median node connectivity score of 28.5, a mean connectivity of 34.7, and a maximum connectivity of 140. The workflow to identify modules and generate the co-expression network was encapsulated in the exp2gcn2 function from the BioNERO package. We selected a signed network with a soft power threshold of 10, and the Pearson correlation method. We computed the Topological Overlap Matrix (TOM), a measure of network connectivity that emphasizes the shared neighbors between protein pairs to enhance the robustness and reliability of the calculated adjacency network. Using hierarchical clustering on the dissimilarity TOM (disTOM), we identified initial modules of co-expressed proteins. The dynamic tree cut method using the cutreeDynamicTree function with maxTreeHeight of 3, minimum module sizes of 35 and deep splitting was applied to the protein dendrogram to define these modules by modifying the exp2gcn2 function. The module eigenproteins (MEs), representing the first principal component for the module, were then calculated across modules with moduleEigengenes. These eigenproteins served as characteristic expression profiles of proteins within a module and were used to assess the interrelation between modules. Similar modules were merged based on the eigenprotein dissimilarity, ensuring that highly correlated modules were combined. To improve biological interpretability and reduce redundancy across modules, we iteratively refined the network by adjusting the module merging threshold and evaluating gene ontology enrichment within candidate module merges. Specifically, we merged modules whose eigenprotein correlation exceeded 0.80, as this threshold reflected a high degree of co-expression and overlapping biological function. This strategy preserved distinct co-expression signals while collapsing highly similar modules that shared enriched biological processes.

We used three types of connectivity metrics to describe how each node/protein in the network related to other nodes/proteins. Total connectivity was calculated by summing the weighted correlations between each protein and all other proteins in the network (kTotal). Intra-modular connectivity (kIN) was calculated by summing the weighted correlations between each protein and all other proteins in the assigned module.

#### RNA co-expression network generation using Weighted Gene Co-expression Network Analysis

We adopted the WGCNA methodology to investigate correlations between gene expression values (log_2_ cpm) of DOX-, DAU-, EPI-, MITO- and VEH-treated iPSC-CMs collected three and 24 hours post-treatment as described for the protein network (Langfelder and Horvath 2008). Network parameters for the RNA network were optimized similarly to the protein network. The lowest power threshold that achieved the best scale-free topology fit was β = 16 (R² = 0.9). Modules with eigengene correlations of 0.7 or greater were merged to reduce redundancy. The dynamic tree cut method using the cutreeDynamicTree function with maxTreeHeight of 2, minimum module sizes of 40 and deep splitting was applied to the gene expression dendrogram to define identified modules. Optimization of the soft power threshold and merge threshold by investigating gene ontologies enriched in modules, led to the merging of modules whose eigengenes had a correlation of 0.70 or greater. kIN and ktotal were calculated for all genes in the network. The RNA network had a mean connectivity of 102, median connectivity of 62.5 and max connectivity of 613.

#### Identification of hub proteins & hub genes

We identified hub proteins and genes that might play central roles in the biological processes represented by each module. We used the get_hubs_gcn function to identify hub proteins/genes within our protein abundance and RNA expression networks. Hubs within modules are defined as the genes/proteins with the highest intra-modular connectivity (kIN) score (top 10%) and the highest Pearson correlation value with the module eigenprotein or eigengene (> 0.7).

#### Module correlation to drug treatment

To assess the impact of drug treatments on the identified modules relative to VEH, the correlation between module eigengenes was calculated for each module, treatment, and time point. For each protein and RNA module, the module eigengene or module eigenprotein was calculated using WGCNA’s moduleEigengenes function. This involved taking the expression data of the genes/proteins belonging to a specific module and computing its first principal component across the biological replicates within each specific experimental condition and time point (e.g., VEH 3h, DOX 3h, etc.). For each module in both the RNA and protein networks, the module eigenvalues of each drug treatment group at a specific timepoint (DOX, DAU, EPI, MITO; n = 6 biological replicates per group) was tested for Pearson correlation (*P* < 0.05) with the module eigenvalues of its time-matched VEH (VEH 3h or VEH 24h; n = 6 biological replicates per group). For the 3-hour DOX condition, one biological replicate was identified as an outlier and removed as previously described, resulting in n = 5 replicates for this specific condition. Positive correlations indicate increased eigenvalues of treatment relative to the VEH, reflecting a generally increased expression/abundance in the treatment group for the tested module. Negative correlations indicate decreased eigenvalues of treatment relative to the VEH, reflecting a generally decreased expression/abundance in the treatment group for the tested module.

#### Linear modelling to identify differentially abundant proteins

We utilized the limma package to fit protein abundances to a linear model across conditions (Ritchie et al. 2015; Molania et al. 2019). The RUV-III corrected abundance matrix from six individuals was quantile normalized using the normalizeBetweenArrays function. Drug treatment (DOX, DAU, EPI, MITO or VEH at either three or 24 hours) was modelled as a fixed effect, whereas individual (IND) was treated as a random effect estimated using the duplicateCorrelation function. The linear model fitting was done using lmFit, which incorporated the block effect from the individuals in the design matrix. This model was then passed through the empirical Bayes moderation in the eBayes function to obtain moderated t-statistics. We defined contrasts in the linear model to compare the differential abundance between DOX, DAU, EPI and MITO at either three or 24 hours against their time-matched VEH conditions such that positive log_2_ fold change values correspond to increased abundance in the drug-treated group in comparison to the time-matched VEH, and negative values correspond to decreased abundance in the drug-treated group in comparison to the time-matched VEH. The model was refitted with these contrasts, and empirical Bayes moderation was applied again to adjust the statistics. This summary was visually explored through a histogram of nominal and Benjamini-Hochberg adjusted *P* values, whereby observation of the distribution and plotting of abundance values across samples led us to determine that an adjusted *P* < 0.05 was an appropriate cutoff for significance for differential abundance. We denote proteins that meet this criterion as differentially abundant proteins (DAPs).

#### Linear modelling to identify differentially expressed genes

We performed pairwise differential expression analysis using the filtered RNA counts matrix and the edgeR-voom-limma pipeline contrasting each treatment against the VEH at each timepoint as described for proteins (Law et al. 2018). Differentially expressed genes (DEGs) are defined as those genes for each drug-VEH pair that meet an adjusted *P* value threshold of < 0.05.

#### Biological process enrichment across modules

To interpret the biological significance of the WGCNA-derived modules, enrichment analysis was performed based on annotated Gene Ontologies (GO) (Aleksander et al. 2023). The enrichGO function from the clusterProfiler R package was used to test the enrichment of terms associated with biological processes in a given set of module proteins against the background of all network proteins (Wu et al. 2021). Likewise, for the RNA network, the enrichGO function from the clusterProfiler R package was used to test the enrichment of terms associated with biological processes in a given set of module genes against the background of all network genes. Enriched terms are those with a Benjamini-Hochberg adjusted *P* < 0.05.

#### Module preservation analysis

To evaluate module preservation between the protein and RNA networks, we utilized the modPres_WGCNA function from the BioNERO package which implements module preservation in WGCNA. When testing for preservation of protein modules, the protein network was designated as the test network and the RNA network was designated as the reference network. Conversely, when testing for preservation of RNA modules, the RNA network was designated as the test network and the protein network was designated as the reference network. In both preservation tests, module preservation was assessed with 1,000 permutations. To assess module preservation, we utilized WGCNA’s Z-summary statistic to quantify various aspects of the correlation structure between networks. Z-summaries > 10 indicated very strong module preservation, suggesting that the gene/protein relationships were highly conserved. Z-summaries between 2 and 10 indicated preservation to a moderate degree, and Z-summaries < 2 indicated that there was no preservation of the module. We designated preserved modules as those having Z-summaries > 2. This resulted in a set of protein modules preserved in the RNA network, and a set of RNA modules preserved in the protein network for further analysis.

#### Protein and RNA module correlation

To compare the overall module expression pattern across treatments, timepoints and phenotypes, we computed Pearson correlation values between module eigengenes (n = 19) from the RNA co-expression network and module eigenproteins from the protein co-expression network (n = 19). Eigengene matrices were first constructed by extracting the first principal component (module eigengene) for each module using WGCNA’s moduleEigengenes function, resulting in matrices where rows correspond to samples and columns to modules. Protein module eigenproteins were aligned with RNA module eigengenes by sample ID, and pairwise Pearson correlations were computed and visualized.

#### Module drug response coherence between RNA and protein

The drug response effect size (log_2_ fold change between each drug and VEH at each timepoint) was obtained from the DAP and DEG analysis. Responses were combined across drugs by calculating the median log_2_ fold change for each protein and gene within their respective networks. We focused on responses at 24 hours given the minimal protein response three hours post-treatment. Each gene shared between the RNA and protein networks (n = 5,231) was associated with a median log_2_ fold change value from the RNA and protein networks. RNA-protein pairs were then grouped as SDR, VDR or NDR based on their protein classification following 24 hours of treatment. We calculated the Pearson correlation in median log_2_ fold change values across RNA and protein.

#### Transcription factor motif enrichment in drug-correlated RNA module promoters

We obtained genomic coordinates for H3K27ac-enriched regions associated with each TOP2i treatment at three and 24 hours post-treatment identified by CUT&Tag from Matthews *et al*. (GEO: GSE291262) (Matthews et al. 2025). Treated iPSC-CMs were collected at the same time for RNA-seq, proteomics and CUT&Tag experiments.

Matthews *et al*. identified 20,137 H3K27ac-enriched regions. These regions were converted to a GRanges object and mapped to the nearest transcription start site (TSS) in human genome (hg38) (Durinck et al. 2009; Lawrence et al. 2013). H3K27ac-enriched regions associated with genes that are expressed in the iPSC-CM network were retained, resulting in 17,163 remaining regions. To focus on the promoter regions of expressed genes, we set a 4 kb window around the TSS of expressed genes that includes 2 kb upstream and 2 kb downstream, and selected H3K27ac-enriched regions overlapping the window by 1 bp. This yielded 5,515 promoter-associated regions. The sequences of the 5,515 promoter-associated regions were obtained using the BSgenome.Hsapiens.UCSC.hg38 package (Team 2023). TF-binding profiles were obtained from the JASPAR2020 database using the JASPAR2020 package in R, and filtered for vertebrate motifs in *Homo sapiens* (Fornes et al. 2020). Motifs were loaded using the getMatrixSet function using the TFBSTools R package (Tan and Lenhard 2016). TF motif enrichment was tested for H3K27ac sequences in RNA-1 and RNA-2 SDR modules against the background of H3K27ac-marked promoter sequences from NDR module genes by using the calcBinnedMotifEnrR function from the monaLisa R package (Machlab et al. 2022). Motifs with an −log_10_(adjusted *P*) > 4 (Fisher’s exact test) were determined to be significantly enriched as per monaLisa R package recommendations.

#### TF enrichment in the RNA and protein networks

TFs associated with the set of tested TF motifs described above were obtained from the JASPAR2020 database and subset to those TFs expressed in the RNA network or detected as protein in the protein network. We tested for enrichment of TF genes in each individual module in either the RNA or protein network against the background of all other modules in the RNA network or protein network. TF-enriched modules are defined as those with a Fisher’s exact test *P* < 0.05 and odds ratio greater than one.

#### Protein and RNA network connectivity of enriched transcription factors

TF genes associated with enriched TF motifs were parsed from TF genes expressed in the RNA network which were not associated with any enriched TF motifs. Intra-modular connectivity (kIN) between TF genes associated with enriched motifs was compared to TF genes that are not associated with enriched TF motifs by Wilcoxon test, where *P* < 0.05 was considered significant. TF connectivity in the protein network was similarly tested.

#### TP53 target enrichment in modules within the RNA and protein networks

TP53 target genes, identified through integration of multiple TP53 ChIP-seq experiments, were obtained from Fischer *et al*. (Fischer 2017) and subset to those genes expressed in the RNA network or detected as protein in the protein network. We tested for enrichment of the TP53 target genes in each individual module in either the RNA or protein network against the background of all other modules in the RNA network or protein network. TP53 target-enriched modules are defined as those with a Fisher’s exact test *P* < 0.05 and odds ratio greater than one.

#### CVD and CF GWAS enrichment testing

We obtained GWAS summary statistics from the GWAS catalog (Cerezo et al. 2025). We downloaded summary statistics for “heart disease” (EFO_0003777) and “heart function measurement” (EFO_0004311) and selected the set of mapped genes as defined by the summary statistics reported by the original study. We first combined the set of mapped genes of traits identified through multi-trait analysis of GWAS (MTAG) with their non-MTAG terms to limit trait redundancy. We then combined the mapped genes of traits from “Anthracycline-induced cardiotoxicity in early breast cancer”, “Anthracycline-induced cardiotoxicity in childhood cancer”, and “Anthracycline-induced cardiotoxicity in breast cancer” into one trait termed “Anthracycline-induced cardiotoxicity” to capture the broadest set of genes specifically for this AC-related trait. This resulted in 1,077 traits to test. We tested for enrichment of cardiovascular function and disease traits within modules of the protein network.

Module-wise enrichment testing was performed by comparing the overlap of risk proteins for a particular trait in each module to the background overlap of risk proteins for the trait with the combined set of all other modules. A Fisher’s exact test was used to determine the statistical significance of the overlap and was repeated across all traits for each module. Enriched traits are defined as those with a Fisher’s exact test adjusted *P* < 0.05 and odds ratio greater than one.

#### Longitudinal strain-associated protein enrichment testing

To assess the enrichment of LS-associated proteins within modules of the protein network, we obtained a set of 703 LS-associated proteins from plasma that were identified from 2,956 individuals with low prevalence of CVD (Perry et al. 2024). Perry *et al*. regressed protein expression levels against LS measurements to identify proteins with significant positive (β > 0) and negative (β < 0) associations with myocardial strain. These two groups were tested for enrichment in each module in the protein network by Fisher’s exact test. Modules enriched for LS-proteins are defined as those with a Fisher’s exact test adjusted *P* < 0.05 and odds ratio greater than one.

#### Eigenprotein correlation to drug-induced calcium dynamics

Protein eigenprotein data and calcium transient data at 24 hours post treatment for VEH, DAU, DOX, EPI and MITO were filtered to retain the three matched individuals present from Matthews *et al*. (Matthews et al. 2024). Both the eigenprotein and calcium transient data were centered and scaled using z-scores to standardize the variables for correlation analysis. For every module, the 24 hour module eigenvalues for all drug treatments were independently tested for correlation with calcium features including rate, rise slope, decay slope and mean amplitude. Correlations with an adjusted *P* < 0.05 were considered significant.

## DATA ACCESS

Mass Spectrometry data are available from the MassIVE database (https://massive.ucsd.edu) under accession number MSV000099030.

## ACKNOWLEDGEMENTS

We thank all members of the Ward Lab for helpful discussions. We thank Dr. Kelly A. Frazer and the University of California San Diego for providing the iPSC lines through the iPSCORE resource. We thank the Mass Spectrometry Facility at the University of Texas Medical Branch for performing the Mass Spectrometry experiments, and the Flow Cytometry and Cell Sorting Core Facility for access to flow cytometers. This work was funded by a Cancer Prevention Research Institute of Texas (https://www.cprit.state.tx.us/) Recruitment of First-Time Faculty Award (RR190110) to M.C.W, and CPRIT grant RP190682 and RP250644 to W.K.R in partial support of the UTMB Mass Spectrometry Facility. O.D.J was supported by a Jeane B. Kempner Predoctoral Fellowship administered through UTMB.

## Author contributions

O.D.J. and M.C.W conceived and designed the study. E.R.M., S.P, J.A.G, and W.K.R performed the experiments and generated the data. O.D.J analyzed the data. O.D.J and M.C.W wrote the manuscript with input from all authors. M.C.W supervised the project.

## Supplemental information

S1 Appendix: Document containing Supplemental Figures 1-10

S2 Appendix: Document containing Supplemental Tables 1-20

S3 Appendix: Protein abundance matrix

## References

Adegunsoye A, Gonzales NM, Gilad Y. 2023. Induced pluripotent stem cells in disease biology and the evidence for their in vitro utility. Annual Review of Genetics 57: 341–360.

Aleksander SA, Balhoff J, Carbon S, Cherry JM, Drabkin HJ, Ebert D, Feuermann M, Gaudet P, Harris NL. 2023. The gene ontology knowledgebase in 2023. Genetics 224: iyad031.

Almeida-Silva F, Venancio TM. 2022. BioNERO: an all-in-one R/Bioconductor package for comprehensive and easy biological network reconstruction. Functional & Integrative Genomics 22: 131–136.

Aries A, Paradis P, Lefebvre C, Schwartz RJ, Nemer M. 2004. Essential role of GATA-4 in cell survival and drug-induced cardiotoxicity. Proceedings of the National Academy of Sciences 101: 6975–6980.

Arzt M, Gao B, Mozneb M, Pohlman S, Cejas RB, Liu Q, Huang F, Yu C, Zhang Y, Fan X. 2023. Protein-encapsulated doxorubicin reduces cardiotoxicity in hiPSC-cardiomyocytes and cardiac spheroids while maintaining anticancer efficacy. Stem cell reports 18: 1913–1924.

Assum I, Krause J, Scheinhardt MO, Müller C, Hammer E, Börschel CS, Völker U, Conradi L, Geelhoed B, Zeller T. 2022. Tissue-specific multi-omics analysis of atrial fibrillation. Nature communications 13: 441.

Avitabile D, Bailey B, Cottage CT, Sundararaman B, Joyo A, McGregor M, Gude N, Truffa S, Zarrabi A, Konstandin M et al. 2011. Nucleolar stress is an early response to myocardial damage involving nucleolar proteins nucleostemin and nucleophosmin. Proc Natl Acad Sci U S A 108: 6145–6150.

Blischak JD, Carbonetto P, Stephens M. 2019. Creating and sharing reproducible research code the workflowr way. F1000Res 8: 1749.

Borges H, Hesse AM, Kraut A, Coute Y, Brun V, Burger T. 2021. Well Plate Maker: a user-friendly randomized block design application to limit batch effects in large-scale biomedical studies. Bioinformatics 37: 2770–2771.

Bostany G, Chen Y, Francisco L, Dai C, Meng Q, Sparks J, Sessions M, Nabell L, Stringer-Reasor E, Khoury K. 2025. Cardiac dysfunction among breast cancer survivors: role of cardiotoxic therapy and cardiovascular risk factors. Journal of Clinical Oncology 43: 32–45.

Burger B, Vaudel M, Barsnes H. 2020. Importance of block randomization when designing proteomics experiments. Journal of proteome research 20: 122–128.

Burridge PW, Li YF, Matsa E, Wu H, Ong S-G, Sharma A, Holmström A, Chang AC, Coronado MJ, Ebert AD. 2016. Human induced pluripotent stem cell–derived cardiomyocytes recapitulate the predilection of breast cancer patients to doxorubicin-induced cardiotoxicity. Nature medicine 22: 547–556.

Camilli M, Cipolla CM, Dent S, Minotti G, Cardinale DM. 2024. Anthracycline Cardiotoxicity in Adult Cancer patients: JACC: CardioOncology State-of-the-art review. Cardio Oncology 6: 655–677.

Canty Jr JM. 2022. Myocardial injury, troponin release, and cardiomyocyte death in brief ischemia, failure, and ventricular remodeling. American Journal of Physiology-Heart and Circulatory Physiology 323: H1–H15.

Cardinale D, Iacopo F, Cipolla CM. 2020. Cardiotoxicity of anthracyclines. Frontiers in cardiovascular medicine 7: 26.

Cerezo M, Sollis E, Ji Y, Lewis E, Abid A, Bircan KO, Hall P, Hayhurst J, John S, Mosaku A et al. 2025. The NHGRI-EBI GWAS Catalog: standards for reusability, sustainability and diversity. Nucleic Acids Res 53: D998–D1005.

Chambers MC, Maclean B, Burke R, Amodei D, Ruderman DL, Neumann S, Gatto L, Fischer B, Pratt B, Egertson J. 2012. A cross-platform toolkit for mass spectrometry and proteomics. Nature biotechnology 30: 918–920.

Chen Y, Lun AT, Smyth GK. 2016. From reads to genes to pathways: differential expression analysis of RNA-Seq experiments using Rsubread and the edgeR quasi-likelihood pipeline. F1000Research 5: 1438.

Cheng M, Lu X, Huang J, Zhang S, Gu D. 2015. Electrocardiographic PR prolongation and atrial fibrillation risk: a meta-analysis of prospective cohort studies. Journal of cardiovascular electrophysiology 26: 36–41.

Cheng S, Keyes MJ, Larson MG, McCabe EL, Newton-Cheh C, Levy D, Benjamin EJ, Vasan RS, Wang TJ. 2009. Long-term outcomes in individuals with prolonged PR interval or first-degree atrioventricular block. Jama 301: 2571–2577.

Demichev V, Messner CB, Vernardis SI, Lilley KS, Ralser M. 2020. DIA-NN: neural networks and interference correction enable deep proteome coverage in high throughput. Nat Methods 17: 41–44.

Denlinger CS, Sanft T, Baker KS, Broderick G, Demark-Wahnefried W, Friedman DL, Goldman M, Hudson M, Khakpour N, King A. 2018. Survivorship, version 2.2018, NCCN clinical practice guidelines in oncology. Journal of the National Comprehensive Cancer Network 16: 1216–1247.

Durinck S, Spellman PT, Birney E, Huber W. 2009. Mapping identifiers for the integration of genomic datasets with the R/Bioconductor package biomaRt. Nature protocols 4: 1184–1191.

Fischer M. 2017. Census and evaluation of p53 target genes. Oncogene 36: 3943–3956.

Fornes O, Castro-Mondragon JA, Khan A, Van der Lee R, Zhang X, Richmond PA, Modi BP, Correard S, Gheorghe M, Baranašić D. 2020. JASPAR 2020: update of the open-access database of transcription factor binding profiles. Nucleic acids research 48: D87–D92.

Fradley MG, Beckie TM, Brown SA, Cheng RK, Dent SF, Nohria A, Patton KK, Singh JP, Olshansky B, Cardiology AHACoC, et al. 2021. Recognition, prevention, and management of arrhythmias and autonomic disorders in cardio-oncology: a scientific statement from the American Heart Association. Circulation 144: e41–e55.

Friedman GD, Cutter GR, Donahue RP, Hughes GH, Hulley SB, Jacobs Jr DR, Liu K, Savage PJ. 1988. CARDIA: study design, recruitment, and some characteristics of the examined subjects. Journal of clinical epidemiology 41: 1105–1116.

Gagnon-Bartsch JA, Speed TP. 2012. Using control genes to correct for unwanted variation in microarray data. Biostatistics 13: 539–552.

Gilchrist SC, Barac A, Ades PA, Alfano CM, Franklin BA, Jones LW, La Gerche A, Ligibel JA, Lopez G, Madan K. 2019. Cardio-oncology rehabilitation to manage cardiovascular outcomes in cancer patients and survivors: a scientific statement from the American Heart Association. Circulation 139: e997–e1012.

Herrmann GK, Russell WK, Garg NJ, Yin YW. 2021. Poly (ADP-ribose) polymerase 1 regulates mitochondrial DNA repair in an NAD-dependent manner. Journal of Biological Chemistry 296.

Jirkovský E, Jirkovská A, Bavlovič-Piskáčková H, Skalická V, Pokorná Z, Karabanovich G, Kollárová-Brázdová P, Kubeš J, Lenčová-Popelová O, Mazurová Y. 2021. Clinically translatable prevention of anthracycline cardiotoxicity by dexrazoxane is mediated by topoisomerase II beta and not metal chelation. Circulation: Heart Failure 14: e008209.

Joglar JA, Chung MK, Armbruster AL, Benjamin EJ, Chyou JY, Cronin EM, Deswal A, Eckhardt LL, Goldberger ZD, Gopinathannair R. 2024. 2023 ACC/AHA/ACCP/HRS guideline for the diagnosis and management of atrial fibrillation: a report of the American College of Cardiology/American Heart Association Joint Committee on Clinical Practice Guidelines. Journal of the American College of Cardiology 83: 109–279.

Johnson EC, Carter EK, Dammer EB, Duong DM, Gerasimov ES, Liu Y, Liu J, Betarbet R, Ping L, Yin L. 2022. Large-scale deep multi-layer analysis of Alzheimer’s disease brain reveals strong proteomic disease-related changes not observed at the RNA level. Nature neuroscience 25: 213–225.

Johnson OD, Paul S, Gutiérrez JA, Russell WK, Ward MC. 2025. DNA damage-associated protein co-expression network in cardiomyocytes informs on tolerance to genetic variation and disease. iScience.

Knowles DA, Burrows CK, Blischak JD, Patterson KM, Serie DJ, Norton N, Ober C, Pritchard JK, Gilad Y. 2018. Determining the genetic basis of anthracycline-cardiotoxicity by molecular response QTL mapping in induced cardiomyocytes. Elife 7: e33480.

Kong AT, Leprevost FV, Avtonomov DM, Mellacheruvu D, Nesvizhskii AI. 2017. MSFragger: ultrafast and comprehensive peptide identification in mass spectrometry–based proteomics. Nature methods 14: 513–520.

Langfelder P, Horvath S. 2008. WGCNA: an R package for weighted correlation network analysis. BMC bioinformatics 9: 1–13.

Langfelder P, Luo R, Oldham MC, Horvath S. 2011. Is my network module preserved and reproducible? PLoS computational biology 7: e1001057.

Law CW, Alhamdoosh M, Su S, Dong X, Tian L, Smyth GK, Ritchie ME. 2018. RNA-seq analysis is easy as 1-2-3 with limma, Glimma and edgeR. F1000Research 5: ISCB Comm J-1408.

Lawrence M, Huber W, Pagès H, Aboyoun P, Carlson M, Gentleman R, Morgan MT, Carey VJ. 2013. Software for computing and annotating genomic ranges. PLoS computational biology 9: e1003118.

Leduc A, Shipkovenska G, Xu Y, Franks A, Slavov N. 2025. Principles of protein abundance regulation across single cells in a mammalian tissue. bioRxiv doi:10.1101/2025.09.17.676955.

Lee MJ, Yaffe MB. 2016. Protein regulation in signal transduction. Cold Spring Harbor perspectives in biology 8: a005918.

Liu C, Shen M, Liu Y, Manhas A, Zhao SR, Zhang M, Belbachir N, Ren L, Zhang JZ, Caudal A. 2024. CRISPRi/a screens in human iPSC-cardiomyocytes identify glycolytic activation as a druggable target for doxorubicin-induced cardiotoxicity. Cell Stem Cell 31: 1760–1776. e1769.

Liu JE, Barac A, Thavendiranathan P, Scherrer-Crosbie M. 2020. Strain imaging in cardio-oncology. Cardio Oncology 2: 677–689.

Liu Y, Beyer A, Aebersold R. 2016. On the dependency of cellular protein levels on mRNA abundance. Cell 165: 535–550.

Lloyd-Jones DM, Lewis CE, Schreiner PJ, Shikany JM, Sidney S, Reis JP. 2021. The coronary artery risk development in young adults (CARDIA) study: JACC focus seminar 8/8. Journal of the American College of Cardiology 78: 260–277.

Machlab D, Burger L, Soneson C, Rijli FM, Schübeler D, Stadler MB. 2022. monaLisa: an R/Bioconductor package for identifying regulatory motifs. Bioinformatics 38: 2624–2625.

Matthews ER, Abodunrin RO, Hurley JD, Paul S, Gutierrez JA, Bogar AR, Ward MC. 2025. Anthracyclines induce global changes in cardiomyocyte chromatin accessibility that overlap with cardiovascular disease loci. PLoS Genet 21: e1011900.

Matthews ER, Johnson OD, Horn KJ, Gutiérrez JA, Powell SR, Ward MC. 2024. Anthracyclines induce cardiotoxicity through a shared gene expression response signature. PLoS genetics 20: e1011164.

McGowan JV, Chung R, Maulik A, Piotrowska I, Walker JM, Yellon DM. 2017. Anthracycline chemotherapy and cardiotoxicity. Cardiovascular drugs and therapy 31: 63–75.

Molania R, Gagnon-Bartsch JA, Dobrovic A, Speed TP. 2019. A new normalization for Nanostring nCounter gene expression data. Nucleic acids research 47: 6073–6083.

Munro V, Kelly V, Messner CB, Kustatscher G. 2024. Cellular control of protein levels: A systems biology perspective. Proteomics 24: 2200220.

Nakamura T, Mizuno S, Ozawa H, Yamasaki Y, Oya S, Morishige S, Aoyama K, Ymaguchi M, Seki R, Mouri F. 2019. A Functional Analysis of SNP on Anthracycline-Induced Cardiotoxicity in a Uniform Genetic Background Using CRISPR/Cas9 Mediated Single-Base-Edited iPSCs. Blood 134: 5748.

Nielsen JB, Thorolfsdottir RB, Fritsche LG, Zhou W, Skov MW, Graham SE, Herron TJ, McCarthy S, Schmidt EM, Sveinbjornsson G. 2018. Biobank-driven genomic discovery yields new insight into atrial fibrillation biology. Nature genetics 50: 1234–1239.

Ntalla I, Weng L-C, Cartwright JH, Hall AW, Sveinbjornsson G, Tucker NR, Choi SH, Chaffin MD, Roselli C, Barnes MR. 2020. Multi-ancestry GWAS of the electrocardiographic PR interval identifies 202 loci underlying cardiac conduction. Nature communications 11: 2542.

Panopoulos AD, D’Antonio M, Benaglio P, Williams R, Hashem SI, Schuldt BM, DeBoever C, Arias AD, Garcia M, Nelson BC et al. 2017. iPSCORE: A Resource of 222 iPSC Lines Enabling Functional Characterization of Genetic Variation across a Variety of Cell Types. Stem Cell Reports 8: 1086–1100.

Park KC, Gaze DC, Collinson PO, Marber MS. 2017. Cardiac troponins: from myocardial infarction to chronic disease. Cardiovascular research 113: 1708–1718.

Perry AS, Amancherla K, Huang X, Lance ML, Farber-Eger E, Gajjar P, Amrute J, Stolze L, Zhao S, Sheng Q. 2024. Clinical-transcriptional prioritization of the circulating proteome in human heart failure. Cell Reports Medicine 5.

Pfeufer A, Van Noord C, Marciante KD, Arking DE, Larson MG, Smith AV, Tarasov KV, Müller M, Sotoodehnia N, Sinner MF. 2010. Genome-wide association study of PR interval. Nature genetics 42: 153–159.

Ritchie ME, Phipson B, Wu D, Hu Y, Law CW, Shi W, Smyth GK. 2015. limma powers differential expression analyses for RNA-sequencing and microarray studies. Nucleic acids research 43: e47–e47.

Roselli C, Chaffin MD, Weng L-C, Aeschbacher S, Ahlberg G, Albert CM, Almgren P, Alonso A, Anderson CD, Aragam KG. 2018. Multi-ethnic genome-wide association study for atrial fibrillation. Nature genetics 50: 1225–1233.

Roselli C, Surakka I, Olesen MS, Sveinbjornsson G, Marston NA, Choi SH, Holm H, Chaffin M, Gudbjartsson D, Hill MC. 2025. Meta-analysis of genome-wide associations and polygenic risk prediction for atrial fibrillation in more than 180,000 cases. Nature genetics: 1–9.

Sada M, Matsushima S, Ikeda M, Ikeda S, Okabe K, Ishikita A, Tadokoro T, Enzan N, Yamamoto T, Miyamoto HD. 2023. IFN-γ-STAT1-ERK pathway mediates protective effects of invariant natural killer T cells against doxorubicin-induced cardiomyocyte death. Basic to Translational Science 8: 992–1007.

Saleme B, Gurtu V, Zhang Y, Kinnaird A, Boukouris AE, Gopal K, Ussher JR, Sutendra G. 2019. Tissue-specific regulation of p53 by PKM2 is redox dependent and provides a therapeutic target for anthracycline-induced cardiotoxicity. Science translational medicine 11: eaau8866.

Searle BC, Swearingen KE, Barnes CA, Schmidt T, Gessulat S, Küster B, Wilhelm M. 2020. Generating high quality libraries for DIA MS with empirically corrected peptide predictions. Nature communications 11: 1548.

Seyfried NT, Dammer EB, Swarup V, Nandakumar D, Duong DM, Yin L, Deng Q, Nguyen T, Hales CM, Wingo T et al. 2017. A Multi-network Approach Identifies Protein-Specific Co-expression in Asymptomatic and Symptomatic Alzheimer’s Disease. Cell Syst 4: 60–72 e64.

Sturgeon KM, Deng L, Bluethmann SM, Zhou S, Trifiletti DM, Jiang C, Kelly SP, Zaorsky NG. 2019. A population-based study of cardiovascular disease mortality risk in US cancer patients. European heart journal 40: 3889–3897.

Swarup V, Chang TS, Duong DM, Dammer EB, Dai J, Lah JJ, Johnson ECB, Seyfried NT, Levey AI, Geschwind DH. 2020. Identification of Conserved Proteomic Networks in Neurodegenerative Dementia. Cell Rep 31: 107807.

Tan G, Lenhard B. 2016. TFBSTools: an R/bioconductor package for transcription factor binding site analysis. Bioinformatics 32: 1555–1556.

Team T. 2023. BSgenome. Hsapiens. UCSC. hg38: full genomic sequences for Homo sapiens (UCSC genome hg38).

Tebbi CK, London WB, Friedman D, Villaluna D, De Alarcon PA, Constine LS, Mendenhall NP, Sposto R, Chauvenet A, Schwartz CL. 2007. Dexrazoxane-associated risk for acute myeloid leukemia/myelodysplastic syndrome and other secondary malignancies in pediatric Hodgkin’s disease. Journal of Clinical Oncology 25: 493–500.

Thavendiranathan P, Poulin F, Lim K-D, Plana JC, Woo A, Marwick TH. 2014. Use of myocardial strain imaging by echocardiography for the early detection of cardiotoxicity in patients during and after cancer chemotherapy: a systematic review. Journal of the American College of Cardiology 63: 2751–2768.

Van Setten J, Brody JA, Jamshidi Y, Swenson BR, Butler AM, Campbell H, Del Greco FM, Evans DS, Gibson Q, Gudbjartsson DF. 2018. PR interval genome-wide association meta-analysis identifies 50 loci associated with atrial and atrioventricular electrical activity. Nature communications 9: 2904.

Vogel C, Marcotte EM. 2012. Insights into the regulation of protein abundance from proteomic and transcriptomic analyses. Nature reviews genetics 13: 227–232.

Wu T, Hu E, Xu S, Chen M, Guo P, Dai Z, Feng T, Zhou L, Tang W, Zhan L. 2021. clusterProfiler 4.0: A universal enrichment tool for interpreting omics data. The innovation 2.

Xiang G, Shi T, Nwaele CO, Xiao H, Liu Y, Wang Q, Zhang J, Zheng Y. 2025. Inhibition of the Sp1/PI3K/AKT signaling pathway exacerbates doxorubicin-induced cardiomyopathy. Biochimica et Biophysica Acta (BBA)-Molecular Cell Research: 119960.

Yan I, Börschel CS, Neumann JT, Sprünker NA, Makarova N, Kontto J, Kuulasmaa K, Salomaa V, Magnussen C, Iacoviello L. 2020. High-sensitivity cardiac troponin I levels and prediction of heart failure: results from the BiomarCaRE consortium. Heart Failure 8: 401–411.

Yang F, Teves SS, Kemp CJ, Henikoff S. 2014. Doxorubicin, DNA torsion, and chromatin dynamics. Biochimica et Biophysica Acta (BBA)-Reviews on Cancer 1845: 84–89.

