## Supplementary material for "Conserved RNA–protein modules link early anthracycline responses to atrial fibrillation risk": S1 Appendix

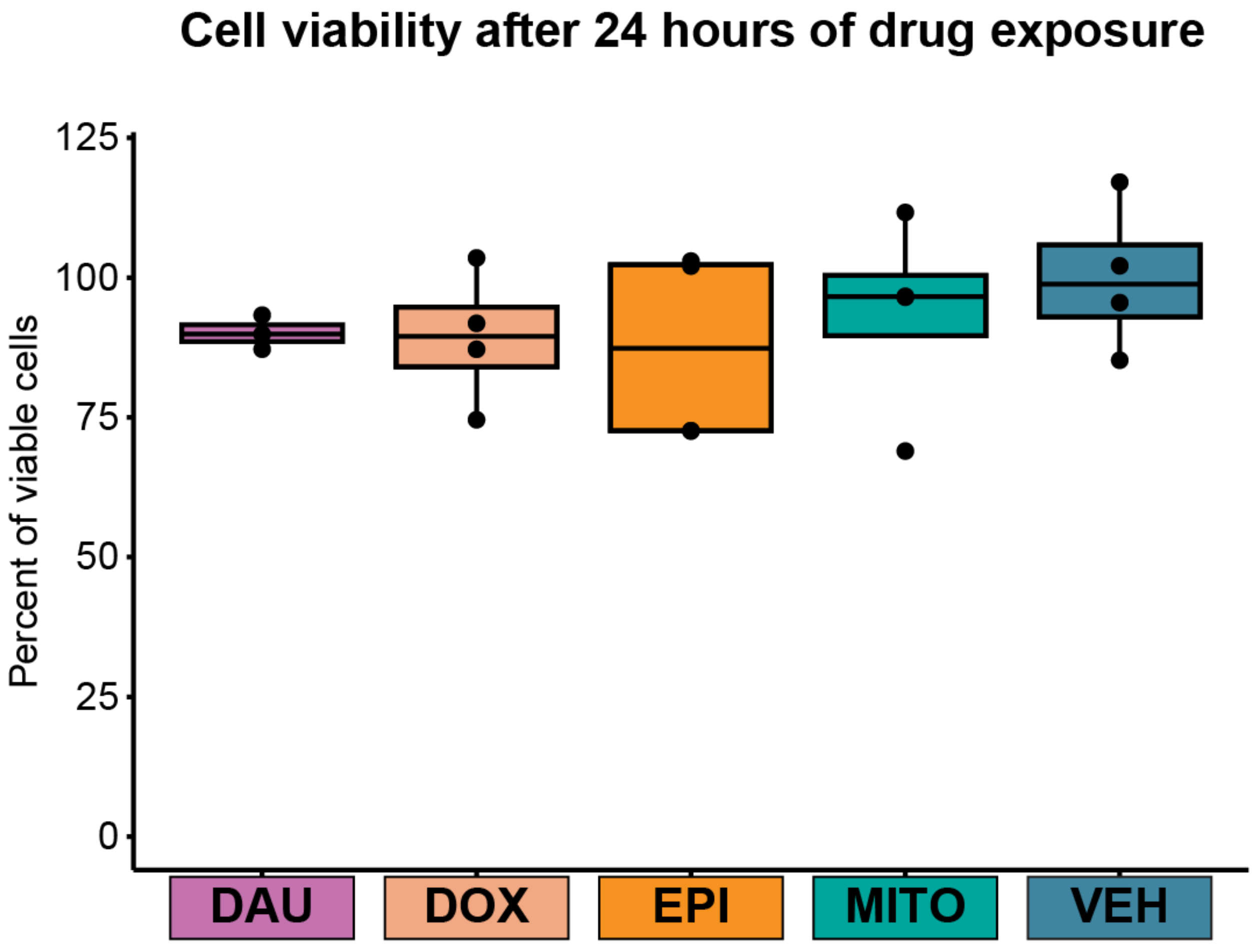


**Figure S1: Treatment of iPSC-CMs with 0.5 µM TOP2i does not affect cell viability.** Individual 5 iPSC-CM viability following 24 hours of treatment with 0.5 µM DAU, DOX, EPI, MITO and VEH normalized to VEH. Viability was measured in quadruplicate for each treatment.

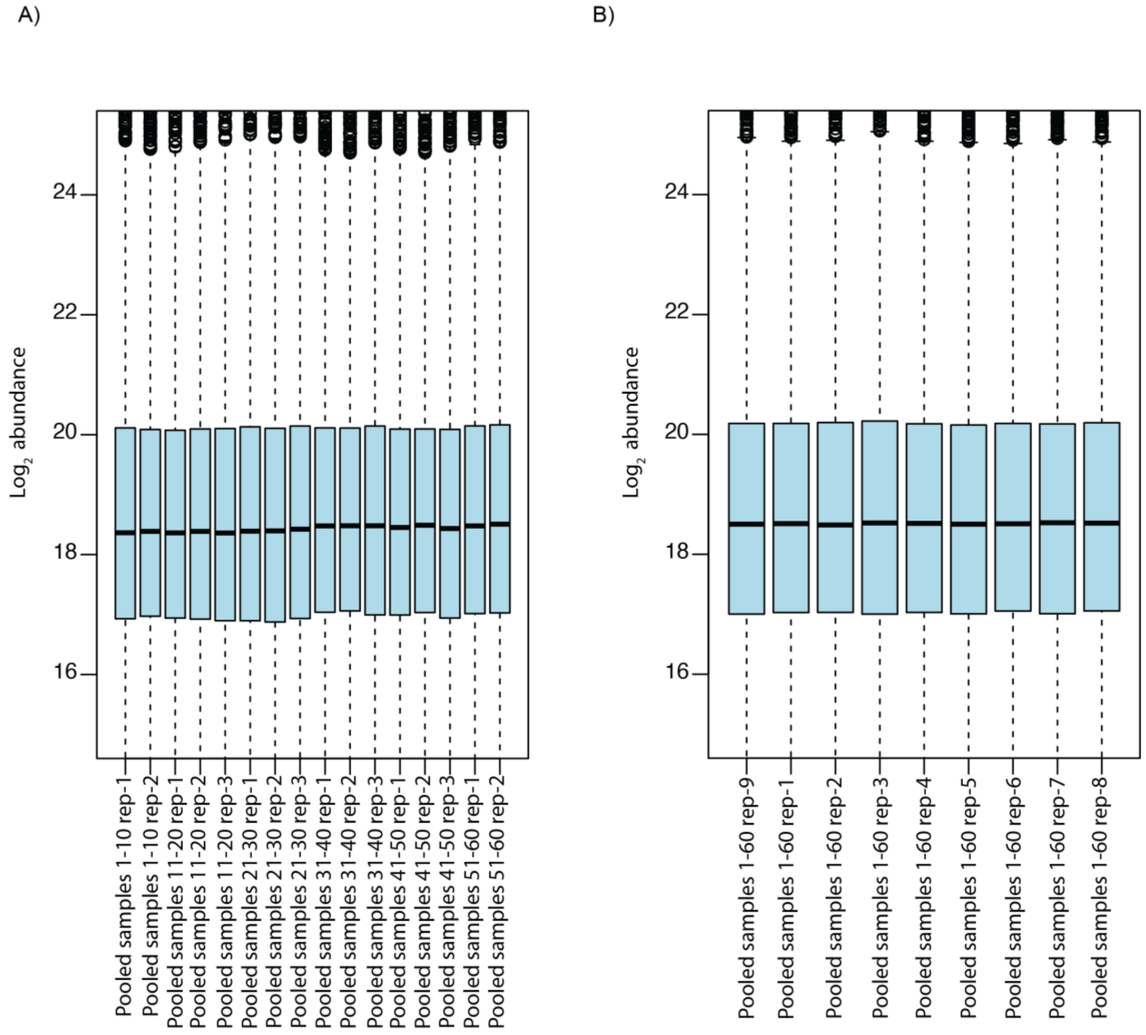


**Figure S2: $Log_2$ protein abundance varies minimally across mass spectrometry runs.** Protein samples were block randomized on the mass spectrometer such that all treatments from an individual comprised a batch. To account for technical variability introduced by the mass spectrometer, samples were pooled either by all treatments from an individual in a batch (n = 10 samples) or pooled across all experimental samples across treatments and individuals (n = 60 samples) and quantified over time. **A)** $Log_2$ protein abundance for all proteins among samples pooled by individual and analyzed at the beginning (rep-1) and end (rep-2) of every mass spectrometry run. In some cases, an additional replicate (rep-3) was added to the run given additional room on the machine autosampler. **B)** $Log_2$ protein abundance for pooled samples containing protein from all 60 experimental samples that were run with each individual batch.

A)

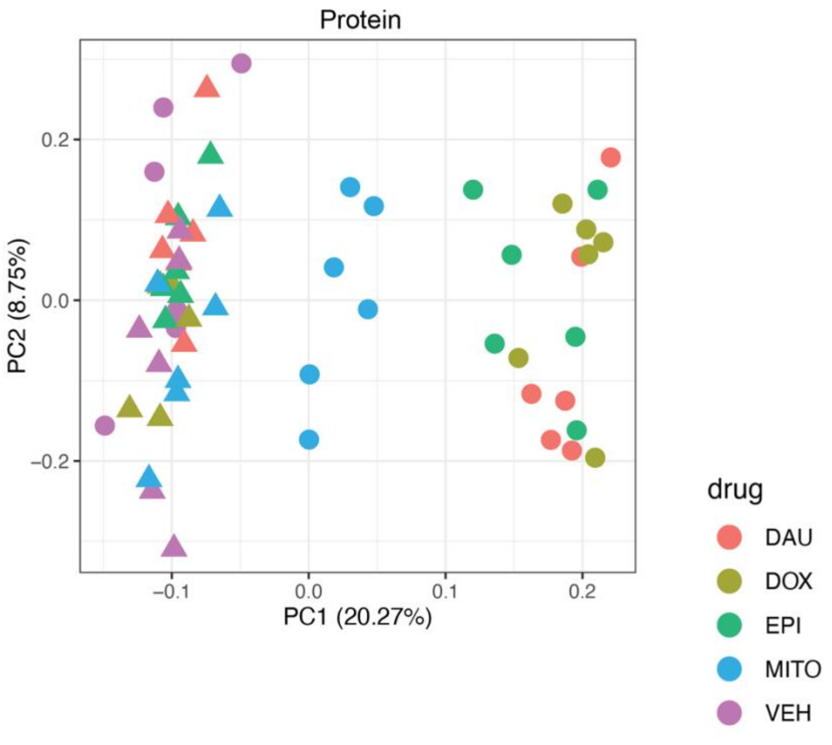


B)

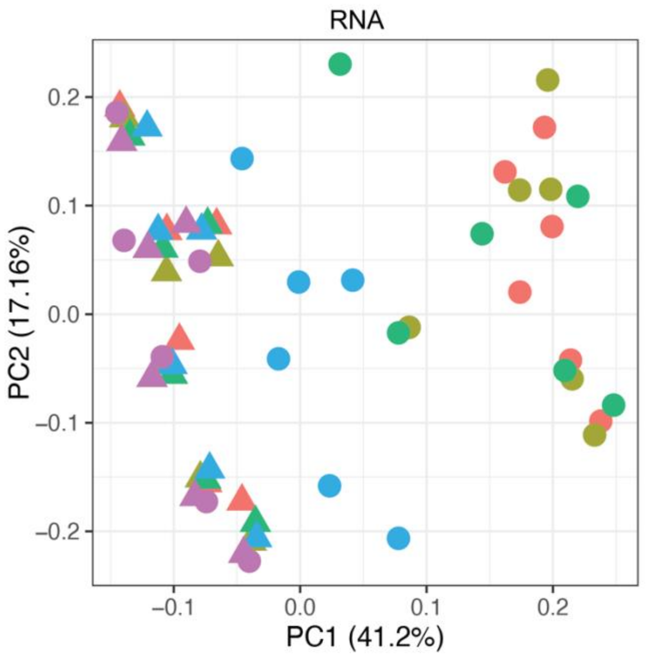


**Figure S3: Treatment duration and drug are the primary contributors to variation in protein abundance and gene expression. A)** Principal component analysis (PCA) of $log_2$ protein abundance data after the removal of unwanted technical variation. Samples are colored by treatment and shaped by treatment duration (Triangle: 3 hours, Circle: 24 hours). **B)** PCA of $log_2$ cpm gene expression data.

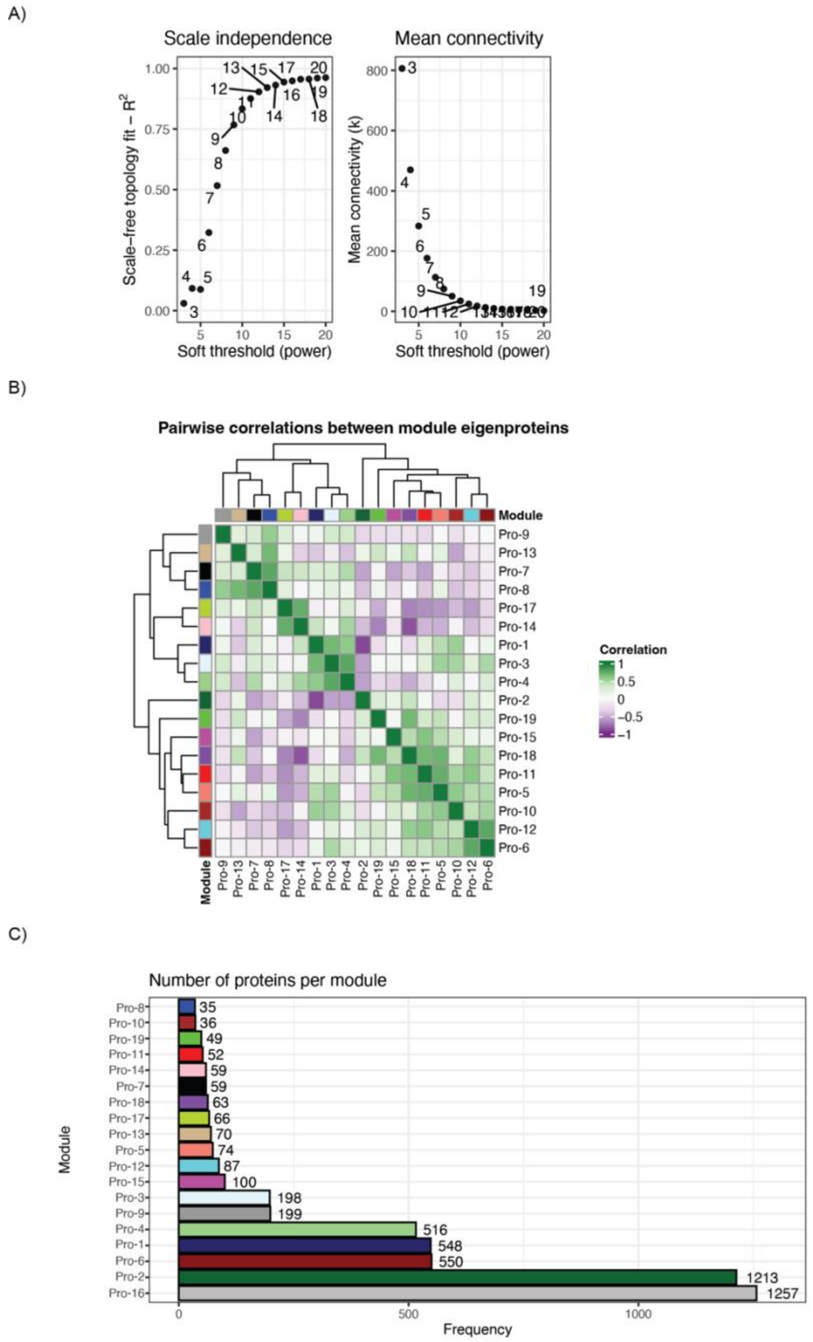


**Figure S4: Weighted protein co-expression network follows a power law and generates co-expressed modules that can be summarized by module eigenproteins. A)** Protein network fit to a scale-free topology across soft power thresholds (left). Fit is determined by the log-log correlation between the connectivity probability P(k) and connectivity (k). Mean network connectivity (k) across soft power thresholds (right). **B)** Correlation heatmap and clustering of module eigenproteins shown in the protein network after merging modules with eigenprotein Pearson correlation greater than 0.8. **C)** Number of proteins in each module in the protein co-abundance network.

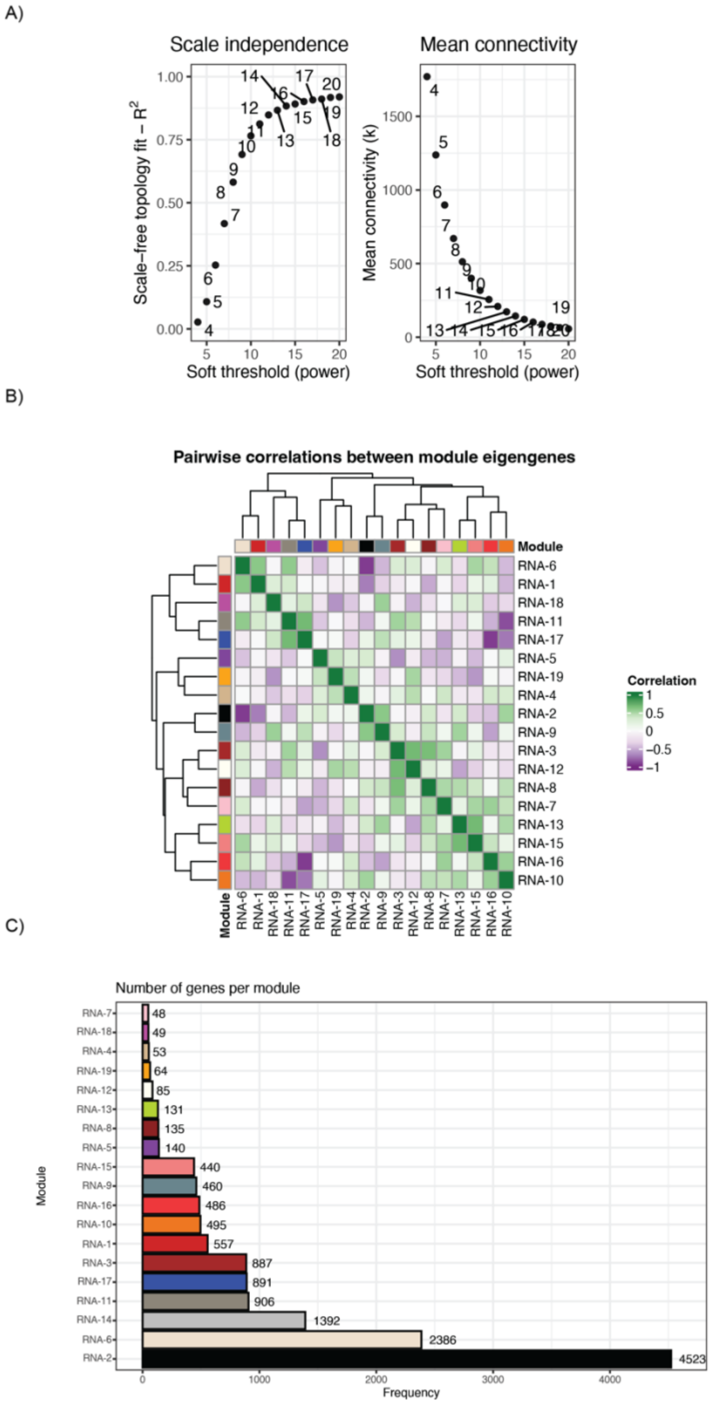


**Figure S5: Weighted gene co-expression network follows a power law and generates co-expressed modules that can be summarized by module eigengenes. A)** RNA network fit to a scale-free topology across soft power thresholds (left). Fit is determined by the log-log correlation between the connectivity probability P(k) and connectivity (k). Mean network connectivity (k) across soft power thresholds (right). **B)** Correlation heatmap and clustering of module eigengenes shown in the RNA network after merging modules with eigengene Pearson correlation greater than 0.7. **C)** Number of genes in each module in gene co-expression network.

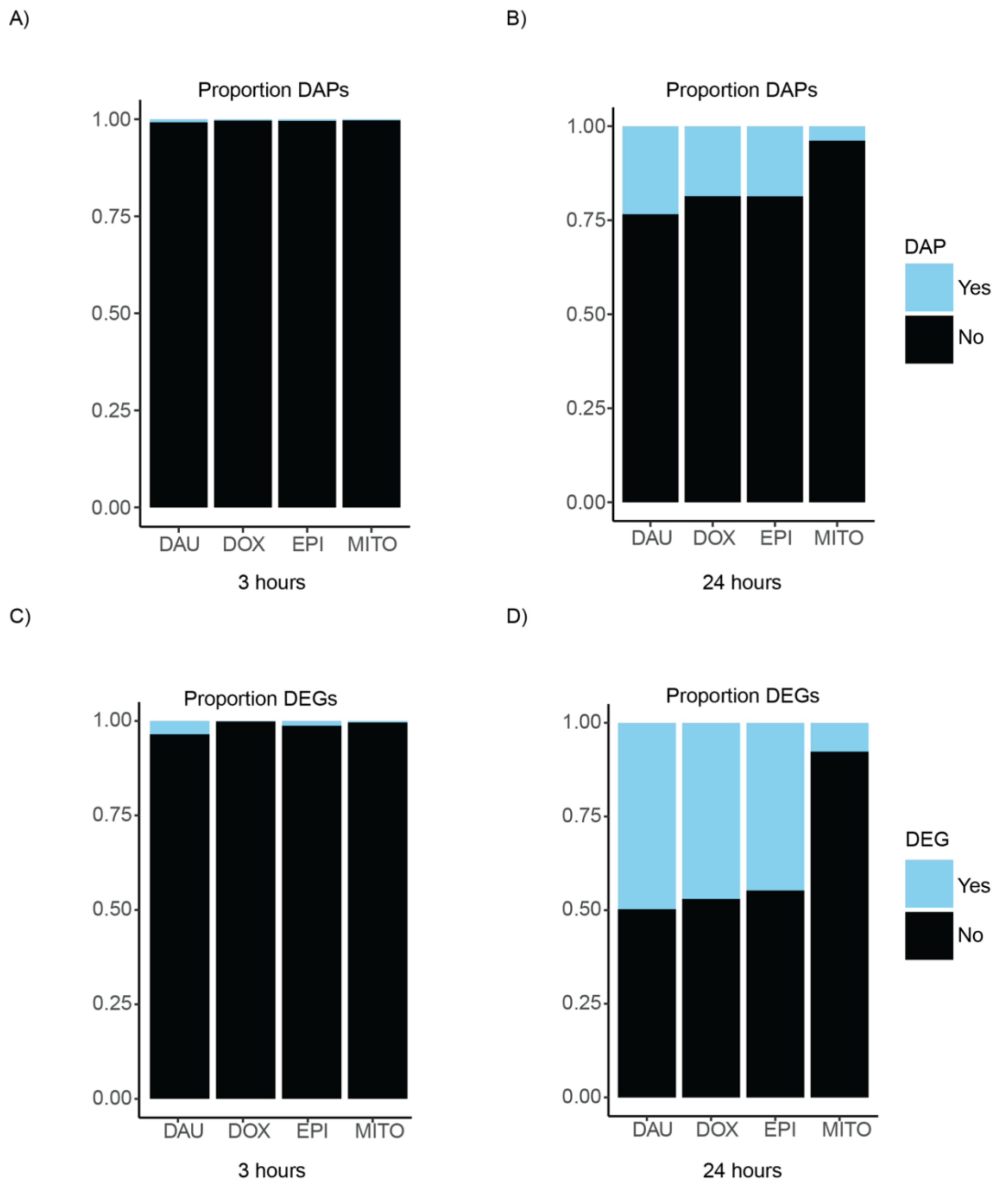


**Figure S6: RNA and protein response to TOP2i is drug- and time-dependent. A)** Proportion of differentially abundant proteins (DAPs) amongst all expressed proteins in each drug comparison for the three-hour timepoint. **B)** Proportion of DAPs in each drug comparison for the 24-hour timepoint. **C)** Proportion of differentially expressed genes (DEGs) amongst all expressed genes in each drug comparison for the three-hour timepoint. **D)** Proportion of DEGs in each drug comparison for the 24-hour timepoint.

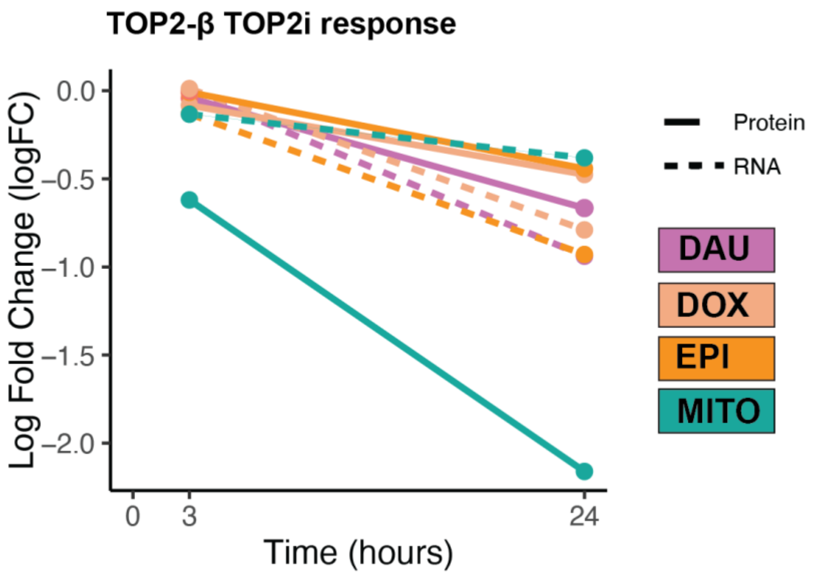


**Figure S7: TOP2β RNA and protein expression is decreased in response to all TOP2i.** TOP2β response to TOP2i ($log_2$ fold change compared to VEH) at the RNA level (dashed line) and protein level (solid line) over time.

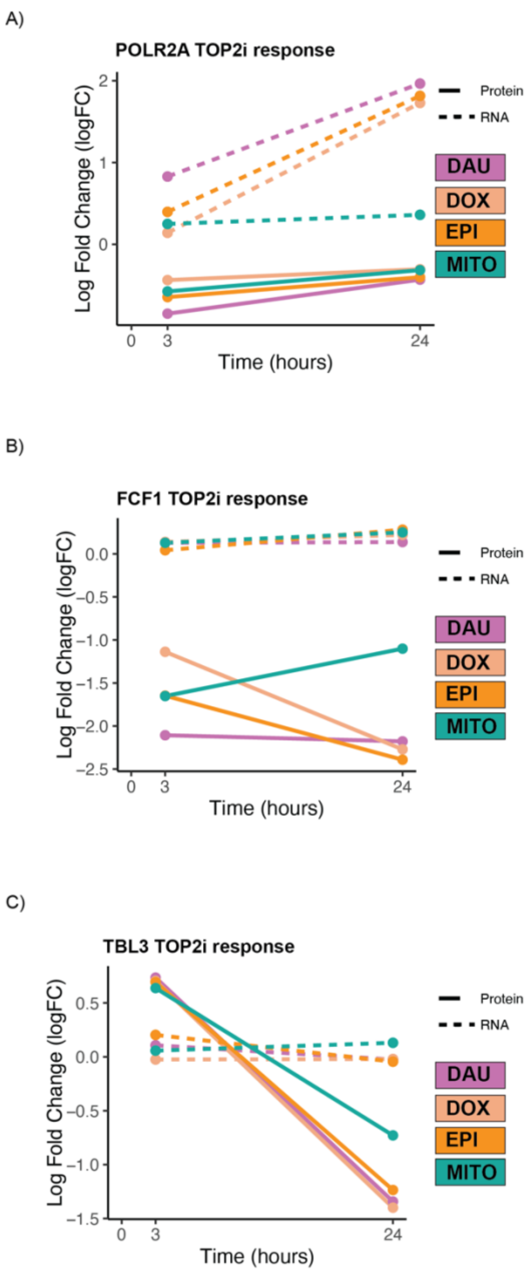


**Figure S8: POLR2A, FCF1 and TBL3 are the three proteins that respond to all TOP2i. A)** POLR2A response to TOP2i ($log_2$ fold change compared to VEH) at the RNA level (dashed line) and protein level (solid line) over time. **B)** FCF1 response to TOP2i. **C)** TBL3 response to TOP2i.

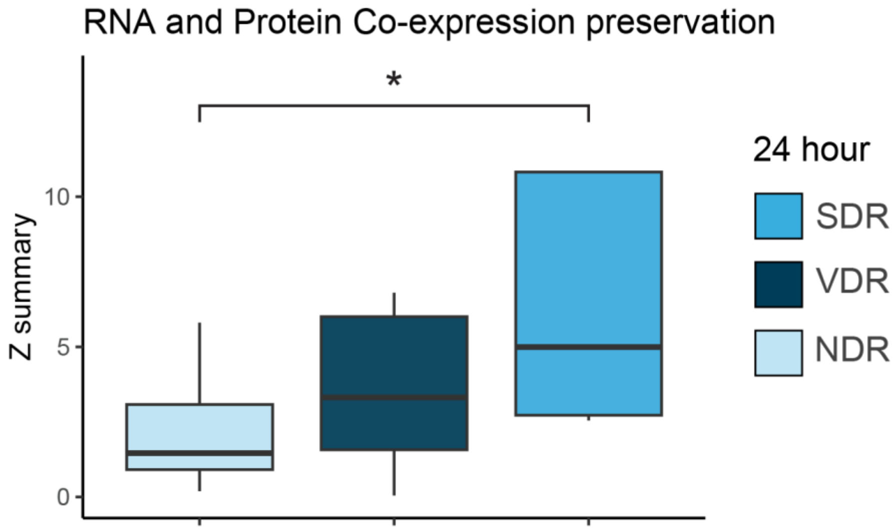


**Figure S9: 24-hour RNA and protein SDR modules are strongly preserved.** Z-summary statistics for RNA and protein preservation were collectively grouped by their 24-hour response to treatment (SDR, VDR or NDR). Z-summary statistics were compared across all groups (*$P$ < 0.05; Wilcoxon ranked sum test).

A)

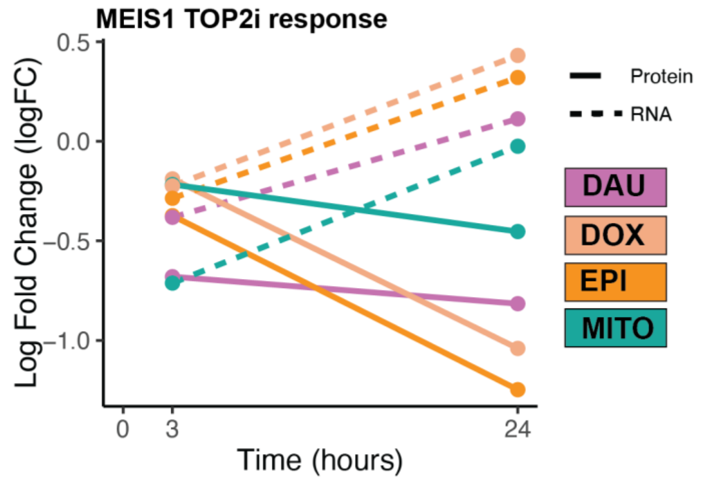


B)

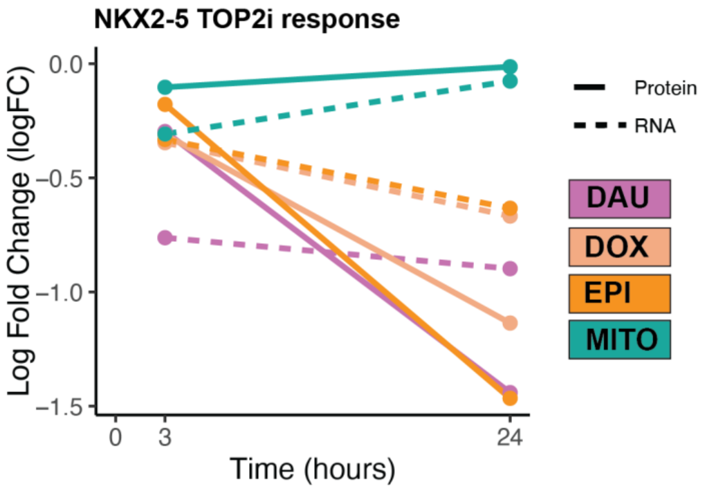


**Figure S10: Transcription factors associated with genetic risk for atrial fibrillation respond to TOP2i treatment. A)** MEIS1 response to TOP2i ($log_2$ fold change compared to VEH) at the RNA level (dashed line) and protein level (solid line) over time. **B)** NKX2-5 response to TOP2i.
